# Cryo-EM Structure of Native Human Uromodulin, a Zona Pellucida Module Polymer

**DOI:** 10.1101/2020.05.28.119206

**Authors:** Alena Stsiapanava, Chenrui Xu, Martina Brunati, Sara Zamora-Caballero, Céline Schaeffer, Ling Han, Marta Carroni, Shigeki Yasumasu, Luca Rampoldi, Bin Wu, Luca Jovine

**Affiliations:** Department of Biosciences and Nutrition, Karolinska Institutet, Blickagången 16, 14183 Huddinge, Sweden.; School of Biological Sciences, Nanyang Technological University, 637551 Singapore, Singapore.; NTU Institute of Structural Biology, Nanyang Technological University, 636921 Singapore, Singapore.; Molecular Genetics of Renal Disorders, Division of Genetics and Cell Biology, IRCCS San Raffaele Scientific Institute, 20132 Milan, Italy.; Cryo-EM Swedish National Facility, SciLifeLab, Tomtebodavägen 23a, 171 65 Solna, Sweden.; Department of Materials and Life Sciences, Faculty of Science and Technology, Sophia University, 7-1 Kioi-cho, Chiyoda-ku, 102-8554 Tokyo, Japan.

## Abstract

Assembly of extracellular filaments and matrices mediating fundamental biological processes such as morphogenesis, hearing, fertilization and antibacterial defense is driven by a ubiquitous polymerization module known as zona pellucida (ZP) “domain”. Despite the conservation of this element from hydra to human, no information is available on the filamentous conformation of any ZP module protein. Here we report the cryo-electron microscopy structure of uromodulin (UMOD)/Tamm-Horsfall protein, the most abundant protein in human urine and an archetypal ZP module-containing molecule, in its mature homopolymeric state. UMOD forms a one-start helix with an unprecedented 180-degree twist between subunits enfolded by interdomain linkers that have completely reorganized as a result of propeptide dissociation. Lateral interaction between filaments in the urine generates sheets exposing a checkerboard of binding sites to capture uropathogenic bacteria, and UMOD-based models of mammalian and avian heteromeric egg coat filaments identify a common sperm-binding region at the interface between subunits.

## INTRODUCTION

The ZP module is a polymer building block of ∼260 amino acids that fold into two topologically related immunoglobulin-like domains, ZP-N and ZP-C (Bokhove and Jovine, 2018; Bork and Sander, 1992; Jovine et al., 2002, 2004). These are separated by a linker that, in crystal structures of non-polymeric precursor forms of ZP module proteins, can be either flexible or rigid, giving rise to different relative arrangements of ZP-N and ZP-C (Bokhove and Jovine, 2018). For example, the precursor of egg coat protein ZP3 is secreted as an antiparallel homodimer where the two moieties of the ZP module are connected by a largely disordered linker, with each ZP-N domain both lying against the ZP-C domain of the same subunit as well as interacting with the ZP-C of the other (Han et al., 2010). On the other hand, the interdomain linker of the UMOD precursor is entirely structured by forming an α-helix (α1) and a β-strand (β1) that pack against ZP-C; this orients the ZP-N domain so that it homodimerizes with ZP-N from another molecule (Bokhove et al., 2016a). Despite these differences, in both cases the last β-strand of ZP-C (βG) - generally referred to as the external hydrophobic patch (EHP) - is part of a polymerization-blocking C-terminal propeptide (CTP) whose protease-dependent release is required for protein incorporation into filaments (Jovine et al., 2004; Schaeffer et al., 2009). Notably, in both mammalian egg coat proteins and UMOD, this process is dependent on membrane anchoring of the precursors (Brunati et al., 2015; Jovine et al., 2002); however, it is unclear how propeptide dissociation triggers polymerization, and the molecular basis of ZP module-mediated protein assembly remains essentially unknown.

To address these questions, we exploited the natural abundance of UMOD (Serafini-Cessi et al., 2003; Tamm and Horsfall, 1950) to study ZP module filaments by cryo-electron microscopy (cryo-EM). First recognized as a major component of hyaline casts in 1873 (Rovida, 1873) and then described as an inhibitor of viral hemagglutination (Serafini-Cessi et al., 2003; Tamm and Horsfall, 1950), UMOD is expressed by cells of the thick ascending limb (TAL) of Henle’s loop as a highly glycosylated, intramolecularly disulfide-bonded and glycophosphatidylinositol (GPI)-anchored precursor. This consists of three epidermal growth factor-like domains (EGF I-III), a cysteine-rich domain (D8C), a fourth EGF domain (EGF IV) and the ZP module, followed by a consensus cleavage site (CCS; often referred to as CFCS in other ZP module proteins) and the EHP-including CTP (Figure S1A) (Bokhove et al., 2016a; Serafini-Cessi et al., 2003). Hepsin protease-mediated cleavage of the CCS (Brunati et al., 2015; Schaeffer et al., 2009) leads to dissociation of mature UMOD from the CTP and triggers its incorporation into homopolymeric filaments that protect against urinary tract infections, reduce nephrolithiasis and are involved in the regulation of water/electrolyte balance and kidney innate immunity (Devuyst et al., 2017; Serafini-Cessi et al., 2003). While common variants of UMOD are strongly associated with risk of chronic kidney disease, higher levels of a monomeric form of UMOD that circulates in the serum and regulates renal and systemic oxidative stress were recently linked to a lower risk for mortality and cardiovascular disease in older adults (LaFavers et al., 2019; Steubl et al., 2019). Thus, elucidating how UMOD polymerization is regulated is not only important for ZP module proteins in general, but also crucial to understand the diverse biological functions of this key urinary molecule.

## RESULTS

### Structure of the UMOD Filament

To obtain information on the supramolecular structure of UMOD, we first imaged human urine samples by cryo-EM. This showed that the protein forms semi-regular sheets through lateral interaction of µm-long filaments, whose pairing generates features that were previously interpreted as the projection of a double-helical structure (Jovine et al., 2002) (Figure 1A). Imaging of purified samples of full-length native UMOD (UMOD_fl_) showed that the majority of filaments had a tree-like structure, resulting from the regular alternation of ∼10 nm-long branches protruding from either side of the polymeric core (Figure 1B); other filaments instead adopted a zig-zag shape consistent with early negative stain EM studies of UMOD (Bayer, 1964) (Figure 1C). In agreement with the observation that the two types of structures occasionally interconvert within individual filaments (Figure S1B), helical reconstruction of UMOD_fl_ showed that these apparently distinct conformations in fact correspond to different views of a single type of filament with 62.5 Å axial rise and 180° twist. The latter parameter, which is even more extreme than the −166.6° helical twist of F-actin (Dominguez and Holmes, 2011), severely complicated structure determination together with the thinness (∼35 Å) and flexibility of the filament core. By averaging 288,403 helical segments, we were however able to solve the structure of UMOD_fl_ to an estimated average resolution of 3.8 Å (Figures 1F-J, S2 and Table S1). To help model building, we also studied native UMOD digested with elastase (UMOD_e_), a protease that removes the entire N-terminal region of the protein (EGF I-III+D8C) by cutting at a single site just before EGF IV (Jovine et al., 2002) (Figure S1A, C and Table S1). This leads to loss of UMOD filament branches (Figures 1D, E and S1D), and comparison of the resulting UMOD_e_ density with that of UMOD_fl_ allowed us to identify the location of the EGF IV N-terminus in the maps (Figures 1F, K and S1E). Using this information, we could unambiguously dock the crystallographic model of UMOD EGF IV and ZP-N (Bokhove et al., 2016a) into the map and then fit the crystal structure of ZP-C (Bokhove et al., 2016a). These placements were validated by the presence of density for the N-glycans attached to ZP-N N396 and ZP-C N513 (Figure 1J, I). Subsequently, a continuous stretch of unexplained density contacting both domains was identified as the ZP-N/ZP-C linker of a third molecule (UMOD 3) that embraces the previously placed ZP-C and ZP-N, which belong to adjacent protein subunits (UMOD 2 and UMOD 4, respectively; Figure 2). This revealed that the relative arrangement of the ZP module moieties of filamentous UMOD is completely different from that of its homodimeric precursor (Bokhove et al., 2016a), so that the distance between the centers-of-mass of the ZP-N and ZP-C β-sandwiches increases from 41 to 91 Å upon polymerization (Figure 3A). Consistent with a ∼120 Å axial periodicity (Jovine et al., 2002), this ZP module conformation allows UMOD monomers to interact head-to-tail (ZP-N-to-ZP-C), with one and two-half subunits per turn (Figures 1F and 2).

**Figure 1.**
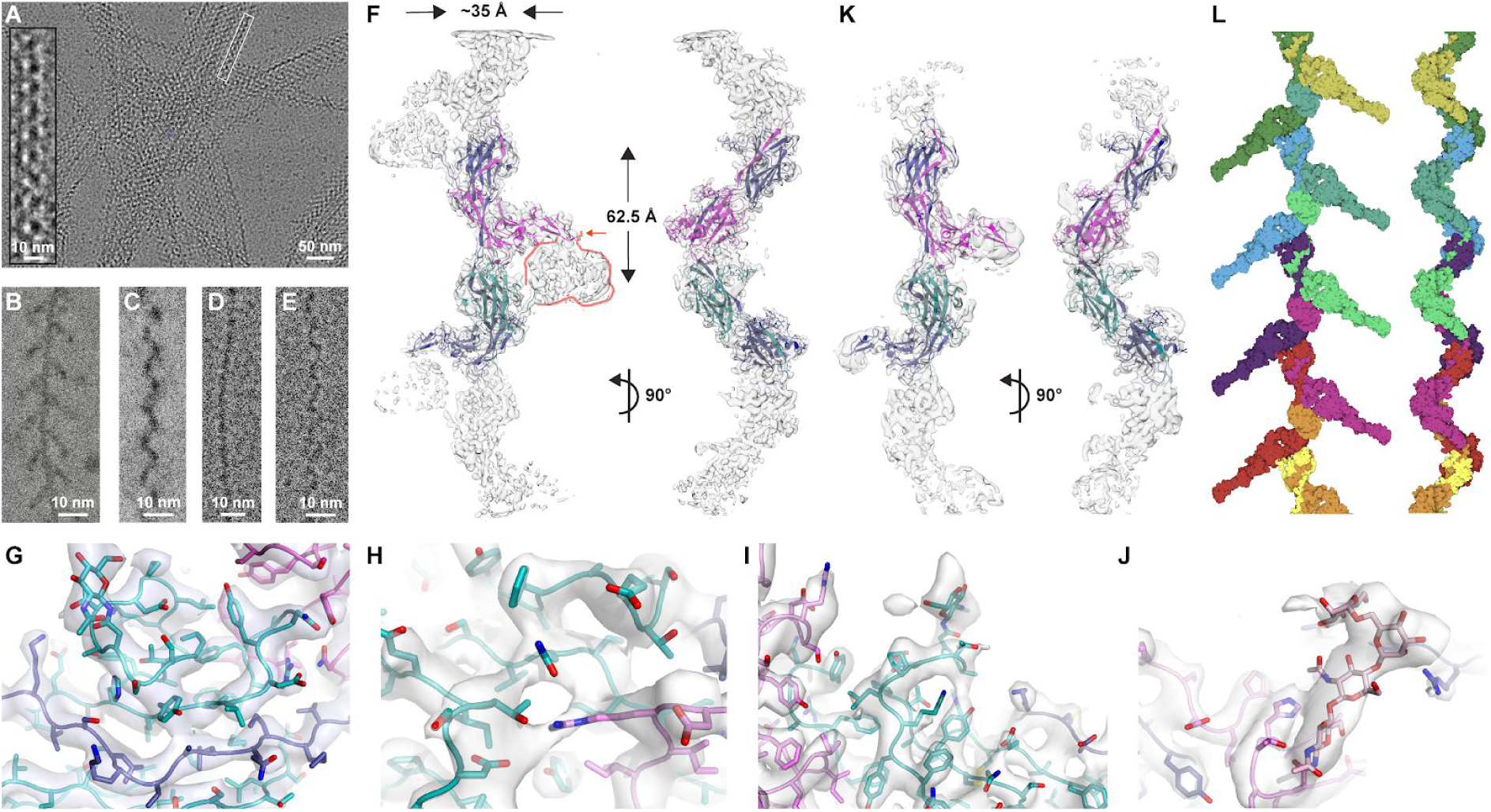
Overall Structure of Human UMOD Filaments. (**A**) Electron micrograph of unstained UMOD filament sheets in human urine. The inset highlights a double helix-like structure resulting from juxtaposition of two individual filaments. (**B, C**) “Tree” front view and “zig-zag” side view of purified native UMOD filaments, imaged using a Volta phase plate. (**D, E**) Front and side views of UMOD_e_ filaments, showing absence of branches. (**F-J**) Orthogonal views of the cryo-EM map of UMOD_fl_ (panel F), oriented as in panels B and C, respectively. Selected sections of the map show the quality of the density for both protein (panels G-J) and carbohydrate residues (panels I-J). The map, which was postprocessed by model-free density modification, is fitted with an atomic model that consists of a complete EGF IV+ZP module (chain A; blue), the ZP-C domain of a second molecule (chain B; cyan) and the EGF IV+ZP-N domain of a third one (chain C; magenta). (**K**) Density-modified cryo-EM map of UMOD_e_, in two orthogonal views oriented as in panels D, E. Comparison of this map with that of UMOD_fl_ identifies density belonging to the N-terminal half of UMOD (salmon contour in the front view of panel F), which is lost upon site-specific cleavage by elastase (orange arrow). (**L**) Goodsell-style depiction of a complete UMOD_fl_ filament model, with protein subunits shown in different colors.

**Figure 2.**
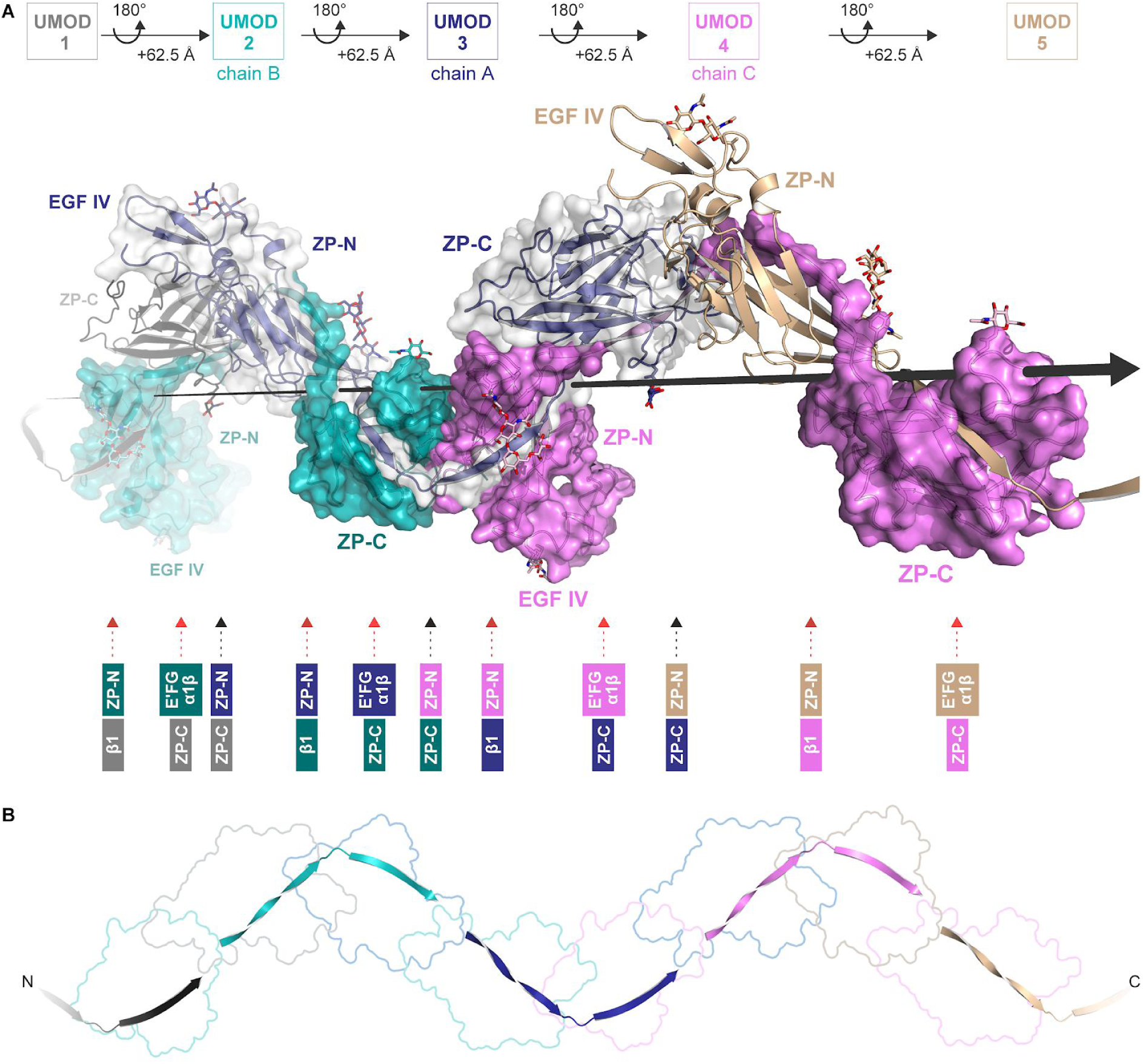
Interaction Between Each Copy of UMOD and the ZP Modules of Four Other Subunits Generates a Unique Filament Architecture. (**A**) A section of a UMOD filament is shown that consists of 5 consecutive subunits (UMOD 1-5) related by the helical symmetry operation indicated in the top panel. In the middle panel, where the helical axis is represented by a large black arrow, subunits are depicted in cartoon (UMOD 1, 3 and 5) and surface (UMOD 2, 3 and 4) representation to highlight protein-protein interfaces (with UMOD 1 ZP-N and UMOD 5 ZP-C omitted for clarity). In the filament, the ZP-N/ZP-C linker of each molecule (for example, UMOD 3) wraps around the ZP-C domain of the subunit that precedes it (UMOD 2) and the ZP-N domain of the subunit that follows it (UMOD 4); additionally, the ZP-N and ZP-C domains of the same molecule are engaged in interactions with the ZP-C domain of the subunit that in turn precedes UMOD 2 (UMOD 1) and the ZP-N domain of the subunit that follows UMOD 4 (UMOD 5), respectively. As summarized in the bottom panel, every UMOD subunit is thus interacting with another four by being engaged in six interfaces that belongs to three different types (ZP-N/ZP-C, black arrow; ZP-N/β1, dark red arrow; E’FG,α1β/ZP-C, light red arrow). Subunits 3, 2 and 4 in this figure correspond to Figure 3 chains A, B and C, respectively. (**B**) Path of the interdomain linkers of UMOD1-5, whose domains are outlined in the background. The view is rotated by ∼40° around the Y axis, compared to panel A.

**Figure 3.**
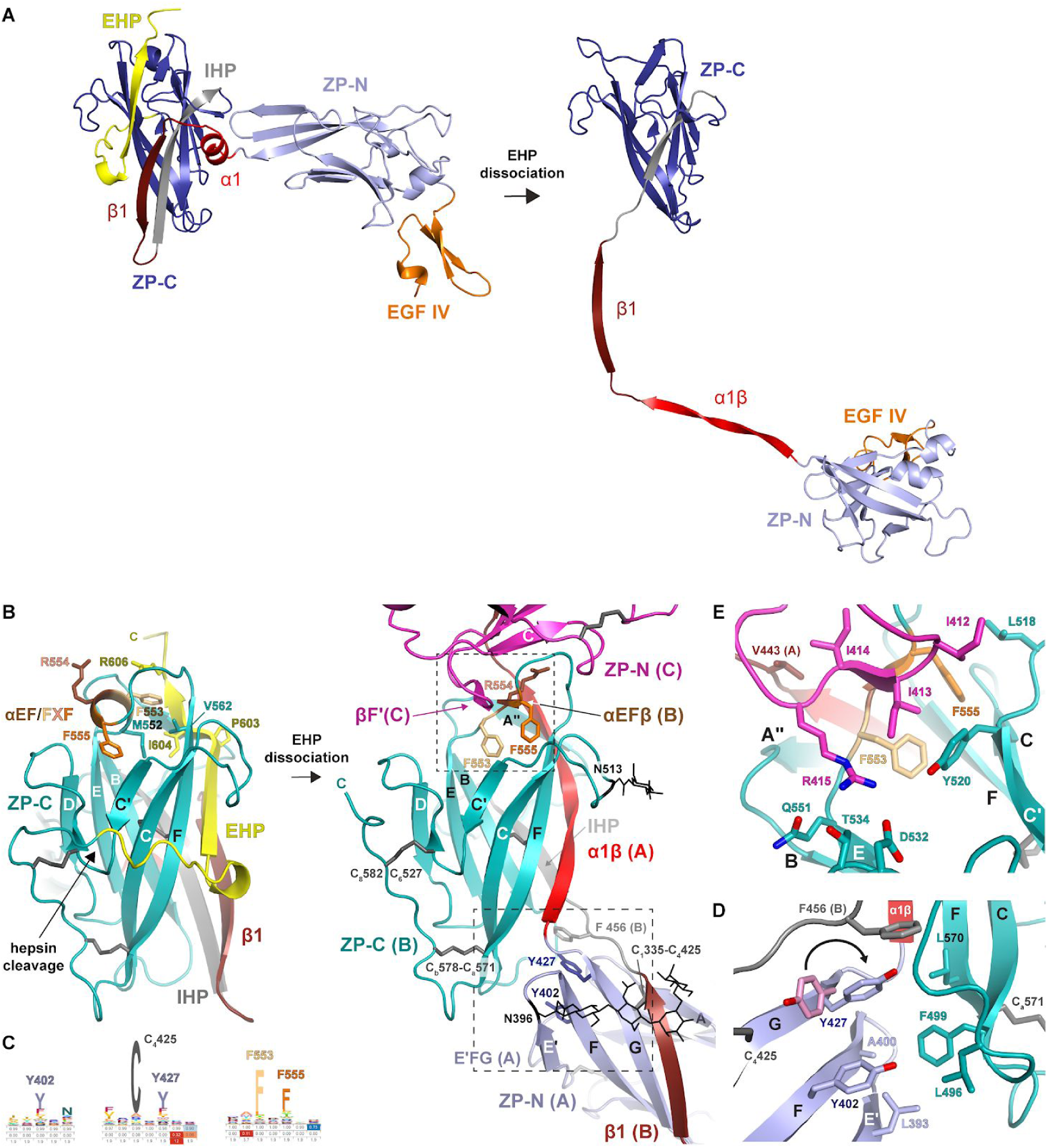
Conformational Changes and Protein-Protein Interactions Underlying UMOD Polymerization. (**A**) Comparison of the precursor and polymeric structures of UMOD shows how dissociation of the EHP triggers a major conformational change of the ZP module. This involves a significant rearrangement of the interdomain linker, which not only completely dissociates from ZP-C but also changes secondary structure upon conversion of α1 in the precursor to α1β in the polymer. Molecules are depicted in cartoon representation, with only one subunit of the UMOD precursor homodimer shown; structural elements are colored as in Figure S1A, with the N- and C-terminal halves of the ZP-N/ZP-C linker colored bright and dark red, respectively. (**B**) In the ZP-C domain of the precursor form of UMOD (left panel, teal), the polymerization-blocking EHP β-strand interacts hydrophobically with a short α-helix (αEF) encompassing the FXF motif. Hepsin-mediated cleavage of the CCS of this molecule (chain B/UMOD 2) triggers release of its EHP, which is replaced by α1β from the interdomain linker of a second UMOD subunit (chain A/UMOD 3) already incorporated at the end of a growing filament (right panel). This allows the FXF motif of molecule B to form an intermolecular β-sheet (αEFβ/βF’; upper dashed box) with the ZP-N fg loop of a third, incoming subunit (chain C/UMOD 4, magenta). Another result of the CCS cleavage is that the C-terminus of mature UMOD 2 is freed for interaction with the D8C domain of the same molecule (not shown). Elements are shown as in panel A, with disulfide bonds and glycan residues represented by thick dark grey and thin black sticks, respectively; β-strands are labelled as in the UMOD precursor (Bokhove et al., 2016a). (**C**) Hidden Markov model logos, highlighting the conservation of selected residues shown in panels B, D and E. (**D**) Hydrophobic interactions stabilize the interface between the E’FG extension of the ZP-N domain of chain A and the ZP-C domain of chain B, corresponding to at the dashed box in the lower right part of panel B. The precursor conformation of ZP-N Y427, which flips during polymerization (arrow) to interact aromatically with IHP F456, is colored pink. (**E**) Details of the interface between the ZP-C domain of chain B and the ZP-N domain of chain C, showing a different view of the area boxed in the upper right part of panel B.

A significant drop in map resolution outside the filament core and lack of close structural homologues precluded accurate modelling of D8C. However, different *ab initio* prediction programs suggested that - consistent with the expected presence of 4 intramolecular disulfides (Hamlin and Fish, 1977; Yang et al., 2004) - this domain adopts a compact fold with average dimensions that closely match the globular density protruding from EGF IV (Figure S3A, B). Notably, the density for the short C-terminal tail of hepsin-processed UMOD, whose flexibility is restricted by the last disulfide of the protein (C_6_527-C_8_582), merges with that of D8C (Figure S3B). This suggests that cleavage of UMOD not only activates its ZP-C domain for polymerization, but also allows it to interact with D8C and, in turn, orient the N-terminal region of the protein relative to the core of the polymer. Although density for the filament branches can only be visualized at low contour levels, suggesting that this part of UMOD is highly flexible, its extent in the map agrees with the dimensions of a homology model of EGF I-III (Figure S3C). By combining the latter with that of D8C and the refined coordinates of EGF IV+ZP, we could assemble a complete filament model consistent with both the tree and zig-zag views of UMOD_fl_ (Figure 1L).

### Filament Formation Involves a Major Conformational Change of the ZP Module’s Interdomain Linker

A dramatic rearrangement of the ZP-N/ZP-C linker region during polymerization underlies the highly different ZP module conformations of the precursor and filamentous forms of UMOD (Figure 3A). In the former, the interdomain linker consists of an α-helix (α1) and a β-strand (β1) that pack against ZP-C β-strand A - an element implicated in polymerization and known as internal hydrophobic patch (IHP; Jovine et al., 2004) - and, in the case of β1, also interact with the EHP (Bokhove et al., 2016a). In polymeric UMOD, α1 and the amino acids that follow it transform into a twisted β-strand that substitutes the EHP of the previous subunit by hydrogen-bonding to its F and A" strands as well as facing βA/IHP (Figure 2A and 3B). Interestingly, an identical copy of the DMKVSL sequence that includes most of α1 (residues 430-435) constitutes β-strand B of UMOD ZP-N (residues 339-344). Antiparallel replacement of EHP/βG by the interdomain linker resembles the donor-strand exchange (DSE) reaction between the subunits of bacterial pili (Waksman, 2017); however, in the case of the ZP module, this β-strand swap is further stabilized by parallel pairing of the linker β1 region to βG of the ZP-N domain of the following subunit. Consistent with the evolutionary conservation of the AG face of the ZP-N domain (Monné et al., 2008), this extends the E’FG β-sheet of the latter, creating a surface against which the well resolved carbohydrate chain attached to N396 packs (Figures 2A and 3B). Because of its interaction with both ZP-C and ZP-N, the interdomain linker of polymeric UMOD acts as a molecular belt that links three consecutive protein subunits, burying a total accessible surface area of 1648 Å^2^ (Figure 2).

### Domain-Domain Interactions in Polymeric UMOD

As a result of the structural changes involving the interdomain linker, the ZP-C domain of one subunit (for example UMOD 2) interacts in different ways with two flanking ZP-N domains that belong to the two subsequent subunits (UMOD 3 and 4) (Figure 2A).

In a first set of contacts, hydrophobic amino acids of UMOD 2 ZP-C, in particular conserved βC F499 and βF L570, interact with residues in the E’FG extension of UMOD 3 ZP-N. These include conserved L393 as well as the signature Tyr of the ZP-N domain (Y402), which acts as a platform for the other near-invariant Tyr of the E’FG extension (Y427) that flips during polymerization in order to stack against highly conserved F456 in the ZP-C IHP (Figure 3B-D). The E’FG extension is another highly conserved feature of the ZP-N fold and distinguishes it from other Ig-like domains (Monné et al., 2008); together with α1 and the IHP, its signature Tyr has long been implicated in ZP module polymerization (Jovine et al., 2004; Monné et al., 2008; Schaeffer et al., 2009).

The second type of interaction, taking place near the C-terminal end of the α1β strand inserted into the ZP-C domain, involves the ef loop of the latter and the fg loop of the ZP-N domain of the subunit following the next (for example, UMOD 2 ef/UMOD 4 fg) (Figure 2A and 3E). Remarkably, the ZP-C ef loop contains the highly conserved FXF motif of the ZP module (553-FRF-555; Figure 3C), which was found to stabilize the homodimeric structure of the precursor of chicken ZP3 (cZP3) by also binding the fg loop of another molecule’s ZP-N. This results in the formation of a short intermolecular β-sheet, involving the FXF motif itself and a hydrophobic sequence of the ZP-N fg loop (139-VII-141) of ZP3 (Han et al., 2010). Although UMOD and ZP3 have different relative domain orientations due to fg loop flexibility and alternative oligomerization states (Figure S4A), the FXF motif of UMOD ZP-C - which in the protein precursor is an α-helix (αEF) - generates an equivalent interface by forming a short β-strand (αEFβ) that pairs with the hydrophobic sequence 412-III-414 in the ZP-N fg loop (Figure 3B). This is followed by highly conserved R415, which inserts into a negatively charged pocket formed by ZP-C Y520, D532, T534 and Q551 (Figure 3E). R415 corresponds to ZP3 ZP-N R142, a residue that binds to ZP-C Y243 and D254 (corresponding to UMOD Y520 and D532, respectively) and is essential for homodimerization and secretion of cZP3 (Han et al., 2010). Notably, the helical conformation of the FXF motif in the precursor form of UMOD is stabilized by the EHP; this immediately suggests how hepsin-dependent dissociation of the latter may facilitate the conformational change that activates ZP-C for interaction with ZP-N from another molecule (Figure 3B).

### Head-to-Tail Mechanism of Polymerization

To complement structural data and functionally investigate the mechanism of filament formation, we first expressed full-length UMOD constructs carrying mutations of the ZP-N fg loop/ZP-C ef loop interface in Madin-Darby Canine Kidney (MDCK) cells, which support UMOD secretion and polymerization (Schaeffer et al., 2009). In agreement with the structure, alanine mutation of ZP-N R415 or a 2-residue deletion affecting the ZP-C FXF motif (ΔF555-A556, ΔFA) do not alter UMOD trafficking but completely abolish its polymerization (Figures 4A and S5A, B). Remarkably, co-transfection experiments showed that, although neither mutant affects the secretion of wild-type UMOD (wt; Figure S5C), ΔFA (but not R415A) has a dominant negative effect on its ability to form filaments (Figure 4B-D). An equivalent result was observed upon co-expression of wt UMOD with a protein variant that cannot polymerize because its CCS has been inactivated (4A (Schaeffer et al., 2009); Figures 4E and S5C). Considering that the dominant negative effect of both the ΔFA and 4A mutations is suppressed in R415A/ΔFA and R415A/4A double mutants (Figure 4F, G), we conclude that - consistent with the structural information (Figure 2) - UMOD is a polar filament whose extension depends on the interaction between an activated ZP-C end and the ZP-N domain of an incoming subunit.

**Figure 4.**
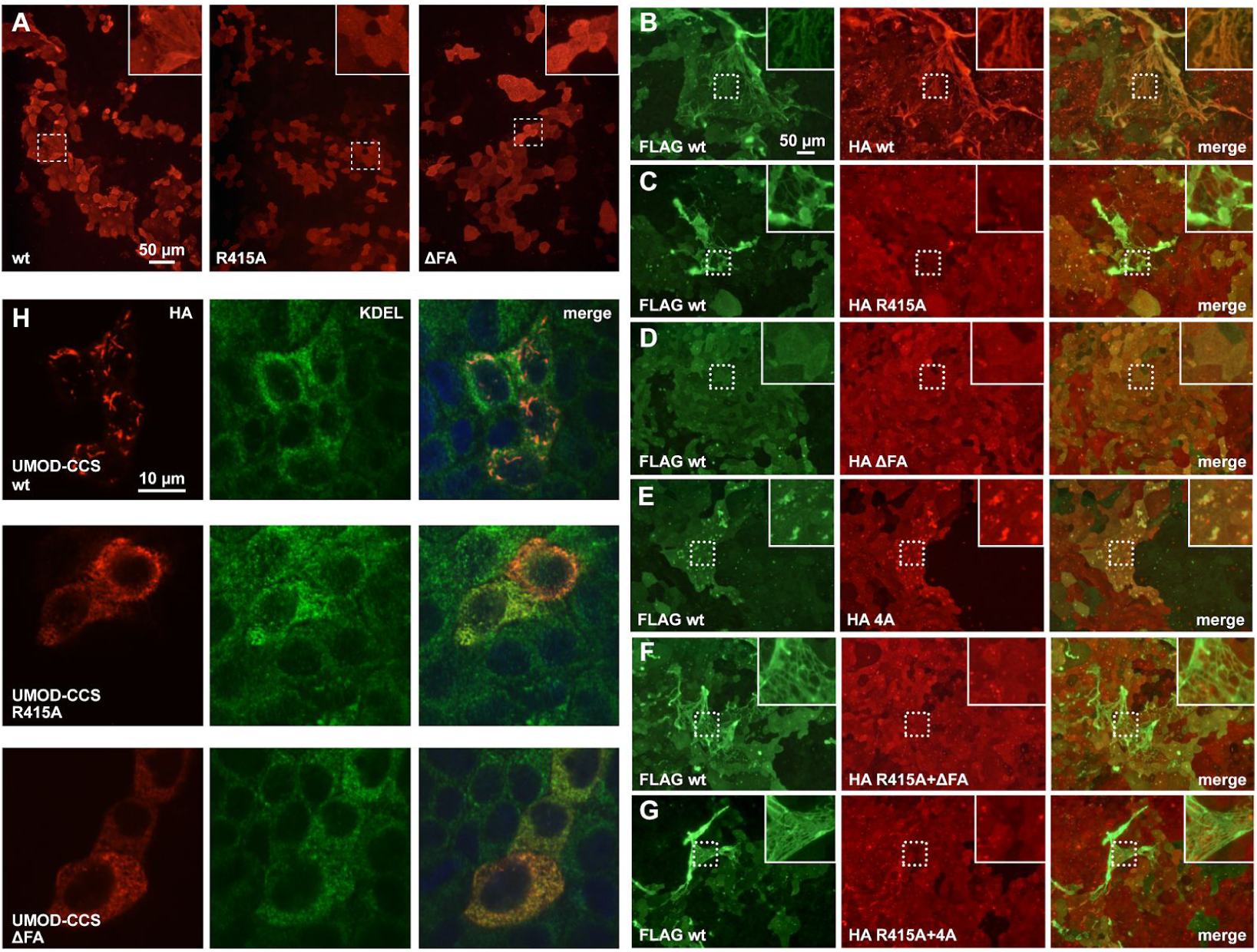
Effect of Polymerization Interface Mutations on UMOD Filament Assembly. (**A**) Immunofluorescence of unpermeabilized stably transfected MDCK cells shows that, unlike wt full-length UMOD, mutants ZP-N (R415A) or ZP-C (ΔFA) do not assemble into filaments. (**B-G**) Immunofluorescence of unpermeabilized cells co-expressing FLAG-tagged wt UMOD (green) and the indicated HA-tagged isoforms (red). UMOD R415A does not incorporate into polymers that only contain wt protein (C). UMOD ΔFA has a dominant negative effect on wt protein polymerization (D) that is rescued in a double mutant carrying both ZP-N and ZP-C mutations (F). Similarly, the dominant negative effect of a CCS mutation (4A) that prevents EHP dissociation (E) is suppressed by the introducing the R415A mutation in the 4A isoform (G). (**H**) Immunofluorescence of permeabilized cells expressing soluble isoforms of UMOD truncated before the EHP shows that wt forms intracellular polymers whereas polymerization interface mutants do not. The intracellular polymers are localized in the endoplasmic reticulum (ER), as shown by co-staining with the KDEL sequence used as ER marker.

A second set of experiments was performed using constructs truncated after the CCS or the EHP. This showed that, unlike UMOD-CCS which is not secreted but forms intracellular polymer-like structures (Schaeffer et al., 2009), UMOD-CCS R415A or ΔFA are impaired in both secretion and polymerization; on the other hand, wt UMOD-EHP and its R415A or ΔFA variants are secreted and do not form intracellular polymers (Figures 4H and S5D, E). These results further confirm the functional importance of the ZP-N fg loop and ZP-C FXF motif. At the same time, they suggest that - although the EHP is ultimately replaced by interdomain linker α1β in the context of the UMOD filament (Figure 3B) - this element is crucial for protein secretion even in the presence of polymerization-impairing mutations. Since UMOD-EHP does also not polymerize extracellularly due to lack of cleavage at the CCS (Figure S5D, F) and wt UMOD only incorporates into filaments attached to the cell from which it was secreted (Figure S5G), this data also supports the idea that EHP dissociation and head-to-tail incorporation into a growing filament are coupled processes occurring at the plasma membrane.

### UMOD Homopolymer Architecture is Conserved in Heteropolymeric Egg Coat Filaments

How similar are the filaments of other ZP module proteins - such as those forming vertebrate egg coats - to the UMOD polymer? The ∼140 Å structural repeat of mouse egg ZP filaments, thought to consist of heterodimers of ZP2 and ZP3 subunits (Wassarman and Mortillo, 1991), closely matches the helical pitch of the UMOD_fl_ polymer (Figure 1F). Moreover, the structure of the latter agrees with the expected solvent exposure of the many glycosylation sites that are variably distributed on the ZP modules of other proteins, including ZP2 and ZP3 (Figure S4B). Similarly, ZP-N- and ZP-C-based superpositions suggest that the N-terminal repeat region of ZP2 and the C-terminal subdomain of ZP3, which have been implicated in sperm binding (Wassarman and Litscher, 2018) but are dispensable for protein incorporation into the growing ZP (Jovine et al., 2002), are compatible with the basic structure of the UMOD filament (Figure S4C).

To gain additional insights into the organization of heteromeric ZP module filaments, we studied native protein complexes solubilized by digesting the unfertilized egg coats (UFE) of medaka fish with high and low choriolytic hatching enzymes (HCE/LCE; Yasumasu et al., 2010). SDS-PAGE and mass spectrometry (MS) analysis of two fractions of this material separated by size-exclusion chromatography (SEC) (F1-2; Figure 5A) revealed that they both contain a ∼37 kDa polypeptide encompassing the ZP module of ZI-3 (the medaka homolog of ZP3), as well as ∼16 and ∼18 kDa fragments corresponding to the ZP-N and ZP-C domains of ZI-1,2 (a subunit that replaces ZP2 in the fish egg coat) (Figure 5B, D, E). Consistent with their different native-PAGE profiles (Figure 5C), native MS of chemically cross-linked F1 and F2 revealed that the latter is a heterocomplex of the three aforementioned species (Figure 5F), whereas the former contains monomers, dimers and trimers of ZI-3 bound to either one or two separate copies of ZI-1,2 ZP-N/ZP-C (Figure 5G). This agrees with the observation that LCE cleaves the interdomain linker of the ZP module of ZI-1,2 (but not that of ZI-3) (Yasumasu et al., 2010), processing a site that corresponds to the short loop between UMOD α1β and β1 (Figure 5H). Moreover, it is consistent with SEC-multiangle light scattering (MALS) evidence that the precursors of ZI-1,2 and ZI-3 are monomeric and monomeric/dimeric, respectively (Figure S6). Taken together, these results strongly suggest that vertebrate egg coat filaments share the same basic architecture as UMOD.

**Figure 5.**
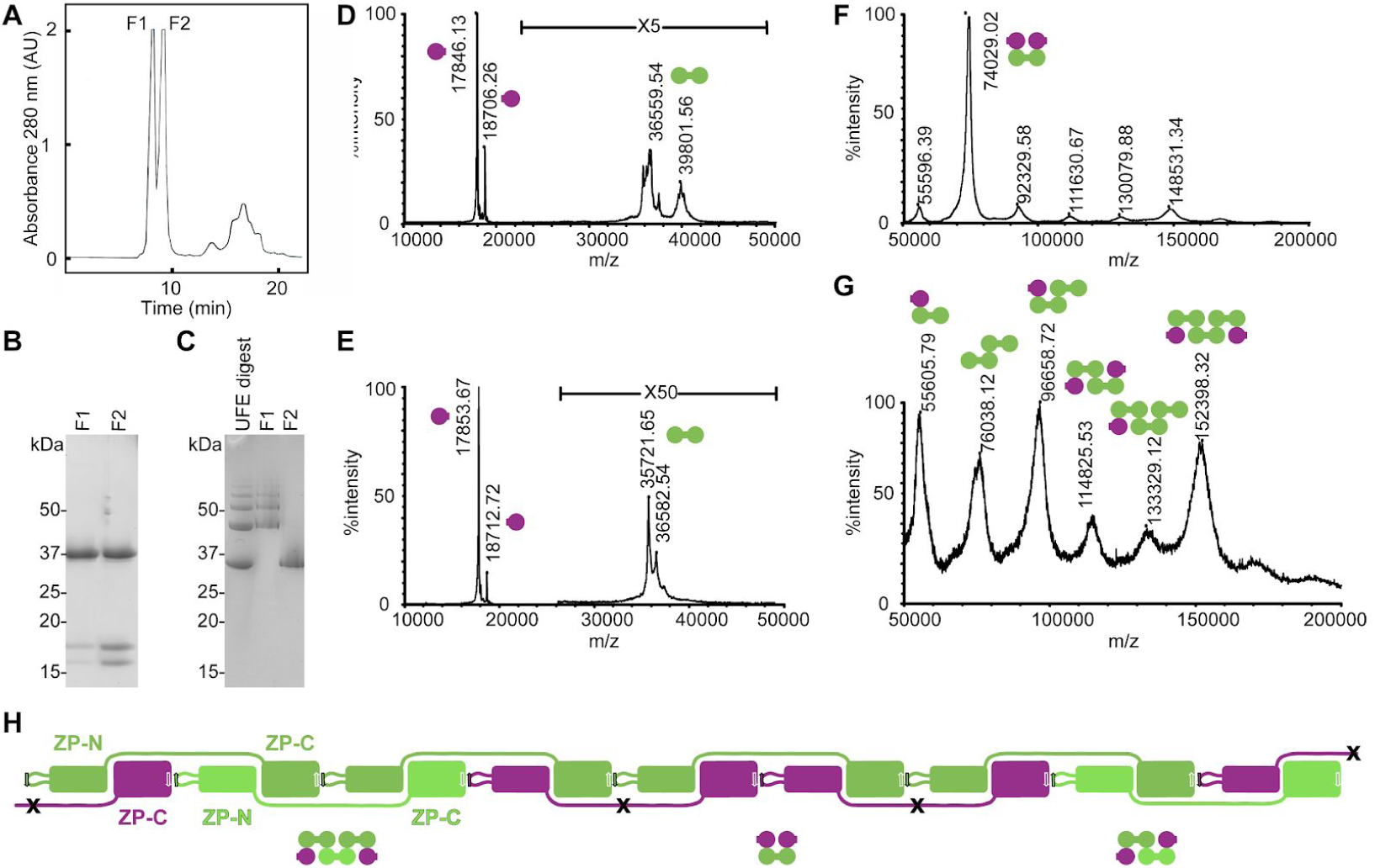
Egg Coat Filament Organization is Consistent with the Architecture of UMOD. (**A**) Analytical SEC of UFE HCE/LCE-digests produces two protein peaks, F1 and F2. (**B, C**) Reducing SDS-PAGE (B) and native-PAGE (C) of SEC purified UFE digests with indicated molecular weight markers. The different SEC elution volumes of F1 and F2 (A) reflect different levels of egg coat filament digestion by HCE/LCE. (**D, E**) TOF-MS spectrograms of purified F2 (D) and F1 (E) products. Round-shaped symbols near the peaks indicate the two moieties of the intact ZP module of ZI-3 (green) or the separated ZP-N and ZP-C domains of LCE-cleaved ZI-1,2 (purple). (**F, G**) TOF-MS spectrograms of cross-linked products of F2 (F) and F1 (G), with domain symbols indicating the deduced subunit composition of cross-linked products. (**H**) Schematic representation of subunit interactions in medaka egg coat filaments. While ZI-1,2 subunits (purple) incorporate into the filaments upon activation of monomeric precursors (Figure S6A), the variable oligomeric state of the ZI-3 precursors (Figure S6B) allows this subunit to be incorporated in either dimeric (green) or monomeric (light green) form. Digestion of the resulting polymers through specific cleavage of the ZI-1,2 interdomain linker by LCE (black crosses) would solubilize filament fragments corresponding to the macromolecular complexes identified in (F) and (G).

## DISCUSSION

The first structure of a polymeric ZP module protein reveals a unique filament architecture whose features, to the best of our knowledge, do not resemble those of any previously reported biological polymer. In particular, the structural information reported in this study clearly shows that, unlike hypothesized in the case of egg coat proteins (Egge et al., 2015; Jovine et al., 2006; Louros et al., 2016; Okumura et al., 2015), UMOD neither polymerizes by forming an amyloid cross-β structure nor assembles through contacts that mostly involve the ZP-N domain or, alternatively, ZP-C. Together with our mutagenesis and biochemical experiments, the structure of filamentous UMOD has long-ranging implications for different aspects of ZP module matrix assembly and function.

### Mechanism of UMOD ZP Module Polymerization

By combining previous crystallographic information on the homodimeric precursor of UMOD (Bokhove et al., 2016a) with the present cryo-EM structure of its filament (Figures 1, 2) and associated mutagenesis experiments (Figures 4 and S5), a four-step mechanism of ZP module-mediated protein polymerization can be proposed (Figure 6). Consistent with the membrane-anchoring requirement for UMOD incorporation into filaments, the process starts when type II transmembrane serine protease hepsin cleaves the CCS of GPI-anchored UMOD precursors (Brunati et al., 2015), which in Figure 6 are represented by two homodimers (UMOD 1 (grey)/UMOD 2 (teal) and UMOD 3 (blue)/UMOD 4 (magenta)) corresponding to the assembled subunits shown in Figure 2. As a result of the orientation of the UMOD homodimer on the membrane and its intrinsic asymmetry (Bokhove et al., 2016a), only one of the moieties of each precursor (UMOD 2 and UMOD 4, respectively) is initially cleaved (step I; Figure 6A). Pulling of the CCS inside the deep specificity pocket of the enzyme, as well as recognition of the substrate’s prime residues (Barré et al., 2014), dislodges the EHP from the corresponding ZP-C domain. This activates the latter for polymerization by releasing its αEF helix and allowing it to engage in an antiparallel β-sheet interaction (αEFβ/βF’) with the fg loop of a ZP-N domain from another dimer (Step II; Figures 3B, E and 6B). Notably, pairing of the ZP-N domains of the homodimeric UMOD precursor forms an extended β-sandwich, oriented so that one ZP-N copy (UMOD 2 and UMOD 4 in Figure 6A, B) sits on top of the other relative to the plasma membrane (Bokhove et al., 2016a). This allows the UMOD branch preceding each ZP-N to project laterally and lie flat on the surface of the membrane; at the same time, it positions the top ZP-N in a favorable position to attack the activated ZP-C of a nearby half-cleaved precursor. Because the top ZP-N copy corresponds to the ZP-C domain that is preferentially cleaved within its own homodimer, interaction between half-cleaved precursors generates a protofilament that remains membrane-bound via the non-cleaved moieties of each homodimer (UMOD 1 and UMOD 3) (Step III; Figure 6C). Importantly, the EHP of the UMOD precursor faces the β1 strand of the interdomain linker, which is paired to ZP-C IHP; detachment of the EHP thus not only affects the αEF helix, but - by loosening the interaction between β1 and ZP-C - also facilitates the conformational change of the interdomain linker and structural transformation of its α1 helix region (Figure 3A and 6B). This is a prerequisite for the final step of assembly, where reorientation of the intact subunit of each incorporated homodimer (UMOD 1 and UMOD 3) as a result of protofilament formation allows it to also be cleaved by hepsin (Figure 6C), locally detaching the nascent polymer from the membrane (Figure 6D). At this point, sequential or concerted conformational changes of the interdomain linkers of the two moieties of each incorporated homodimer allows the newly cleaved subunit (for example UMOD 3) to wrap around the other (UMOD 4), replacing the UMOD 4 ZP-N/UMOD 3 ZP-N interface of the precursor with the UMOD 4 ZP-N/UMOD 3 β1 interaction observed in the filament (Figure 2). This frees the ZP-N of UMOD 3 so that it can engage in its own αEFβ/βF’ interaction with the activated ZP-C domain of UMOD 1 (Figure 2A). At the same time, reminiscent of DSE between the subunits of bacterial pili (Waksman, 2017), the α1β linker region of UMOD 3 fills the open G strand/EHP groove of the activated UMOD 2 ZP-C, facing its IHP (Figure 3B); similarly, the activated ZP-C of UMOD 3 itself is completed by α1β of the UMOD 4 interdomain linker. In both cases, these interactions are stabilized by packing of the ZP-N E’FG extension of one subunit against the ZP-C FC end of the previous (Figure 3B, D).

**Figure 6.**
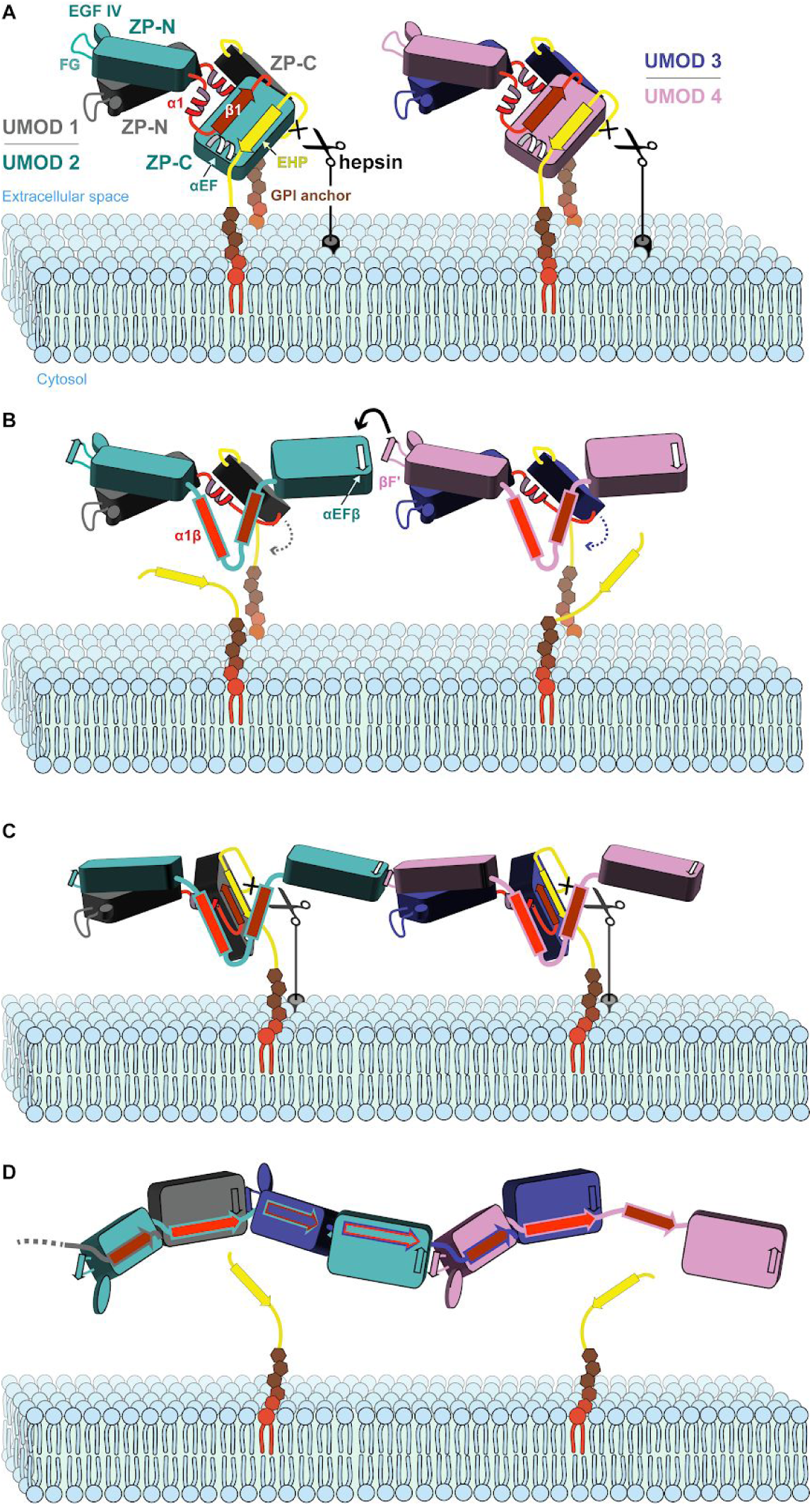
Proposed Mechanism of UMOD Polymerization. (**A**) Hepsin cleaves the membrane-proximal CCS sequence of GPI-anchored UMOD homodimers, triggering the dissociation of the EHP from the corresponding ZP-C domain. Flor clarity, UMOD branch domains preceding EGF IV have been omitted. (**B**) EHP displacement activates ZP-C for polymerization by allowing it to form an intermolecular β-sheet with an incoming ZP-N domain from another homodimer. (**C**) Reorientation of the second ZP-C allows it to also be processed by hepsin, locally detaching the growing filament from the membrane. (**D**) The interdomain linker, whose ordered precursor structure is also perturbed upon EHP dissociation, undergoes a major rearrangement that both replaces the ZP-N/ZP-N interface of the UMOD precursor and compensates the loss of the EHP by DSE, completing filament formation.

The proposed head-to-tail mechanism explains the dominant negative effect of the ΔFA mutation (Figure 4) and, as discussed below, pathologic mutations in human ZP module proteins. By postulating that the protofilament remains membrane anchored, it also rationalizes the observation that activated UMOD subunits do not incorporate into polymers growing on the surface of nearby cells (Figure S5G). Importantly, this provides a solution to the physical problem of assembling the long UMOD polymer in an extracellular environment that is constantly subjected to high flow; clarifies why the growing mammalian ZP thickens from the inside, a process that also depends on membrane anchoring of ZP2 and ZP3 (Jovine et al., 2002; Qi et al., 2002); and is compatible with the recent hypothesis that membrane tethering of ZP module protein α-tectorin is essential for generating layers of extracellular matrix whose progressive release generates the tectorial membrane (Kim et al., 2019a).

Conservation of the structural elements underlying the mechanism, including the length of the interdomain linkers (∼22-26 residues in ZP1-3 compared to 24 residues in UMOD), suggest that other members of the ZP module protein family use a similar mechanism to assemble into filaments that share a common basic architecture. At the same time, the fact that ZP3 precursors can already contain cross-subunit αEFβ/βF’ interactions (Han et al., 2010) may facilitate their incorporation as dimers in the egg coat of species such as chicken (Han et al., 2010), human (Zhao et al., 2004) and fish (Figure 5). In these and other heteropolymeric systems, the variable sequences of the interdomain linkers of different components are also expected to play a major role in determining how subunits incorporate into filaments (Suzuki et al., 2015).

### Functional Implications: From Antibacterial Defense to Fertilization

The structure of polymeric UMOD provides an essential framework to help understanding the biology of this important urinary protein (Devuyst et al., 2017; Serafini-Cessi et al., 2003), as well as all other members of the large superfamily of ZP module-containing extracellular molecules (Litscher et al., 2015). In particular, modelling of UMOD sheets (Figure 1A) by simple juxtaposition of individual filaments (Figures 1L) not only readily explains the apparent double-helical features of the protein’s polymers (inset of Figure 1A and Figure 7A), but also immediately suggests how further supramolecular organization of the latter allows them to trap uropathogenic *E. coli* (UPEC). This is because lateral pairing of multiple filaments generates a surface whose faces expose checkerboard-like arrays of EGF I-III+D8C regions (Figure 7B), each carrying a copy of the high-mannose glycan recognized by type I pilus adhesin FimH (Pak et al., 2001; van Rooijen et al., 1999). A similar strategy is likely to be employed by the major receptor for FimH-positive bacteria in the gastrointestinal tract, glycoprotein 2 (GP2) (Hase et al., 2009; Kolenda et al., 2018), to counteract infection by type-I-piliated *Escherichia and Salmonella* strains. This is because, in agreement with the idea that their genes originated by duplication, GP2 is structurally very similar to UMOD by containing a D8C domain N-terminal to its ZP module (Kobayashi et al., 2004; Yang et al., 2004).

**Figure 7.**
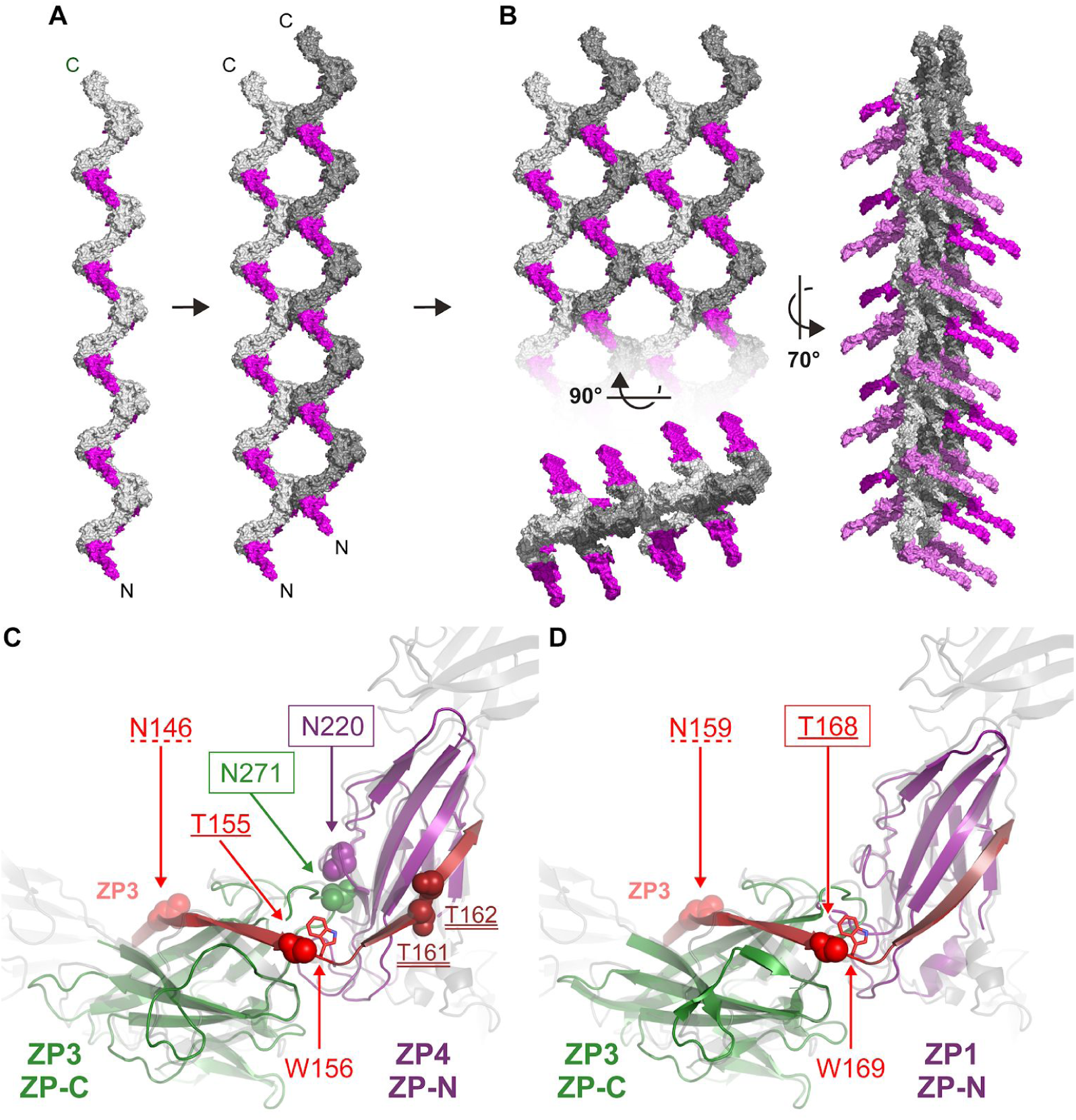
ZP Module Filament Binding Sites for UPEC and Sperm. (A) After assembling as described in Figure 6 and detaching from the plasma membrane of TAL cells, pairs of UMOD filaments aligned parallel with each other can interact laterally to generates structures whose zig-zag view resembles the projection of a double helix (see inset of Figure 1A). The N-terminal branch and C-terminal polymerization regions of UMOD subunits are colored magenta and grey, respectively. (B) Further lateral extension of the filament pairs creates soluble sheets that expose a regular array of UMOD branches in the urine. By carrying the carbohydrate chain that is specifically recognized by the mannose-sensitive fimbriae of UPEC, these surfaces act as double-sided molecular Velcros that capture the pathogens and facilitate their excretion from the body. (**C**) Detail of a porcine ZP filament model, generated by superposition of homology models of adjacent ZP3 and ZP4 subunits (green and purple, respectively) onto the structure of UMOD_fl_ (semi-transparent grey), and projection of the residues of the interdomain linker of another ZP3 subunit onto the UMOD ZP-N/ZP-C linker (red). A cluster of experimentally verified glycosylation sites is shown, with side chains atoms depicted as spheres and boxes highlighting the proximity of ZP3 N271 and ZP4 N220, whose N-glycans mediate sperm binding in pig (Yonezawa, 2014). Note how these residues are also close to the conserved N- (dashed underline) and O- (site 1, single underline; site 2, double underline) glycosylation sites in the ZP3 interdomain linker. The invariant ZP3 Trp that follows site 1 is shown in stick representation at the interface with ZP4. (**D**) Detail of the ZP3/ZP1 interface of an avian egg coat filament model, assembled and represented as described for panel C. The box highlights the location of O-glycosylation site 1, important for *in vitro* binding of chicken ZP3 to sperm (Han et al., 2010).

In yet another part of the body, exposure of functionally important regions of ZP2 and ZP3 subunits adjacent to each other within mammalian egg coat filaments could allow them to cooperate in mediating the initial attachment of gametes at fertilization. Interestingly, modelling of the N-terminal ZP-N repeat region of mammalian ZP2 with the same orientation relative to the filament axis as UMOD’s branch suggest that the second repeat of ZP2 - whose post-fertilization cleavage by cortical granule protease ovastacin is crucial to establish a definitive block to polyspermy (Burkart et al., 2012) - could in principle contact the conserved ZP-C subdomain of ZP3 (Figure S4C). This potential interaction might affect sperm binding by influencing the efficiency of ZP2 cleavage and, like in the case of the D8C domain of UMOD and its C-terminus (Figures 3B and S3B), could depend on the C-terminus of mature ZP3 being freed upon CFCS processing and incorporation into a filament. The ZP-C subdomain of ZP3 could also function by affecting the presentation of the sperm-binding domain of ZP2 or, alternatively, the physical proximity of the subunits may allow them to generate a hybrid sperm-recognition surface that includes regions from both proteins.

Mapping of egg coat protein sites implicated in sperm recognition in mammal and bird onto UMOD-based models of the respective filaments strongly supports the latter concept, which also involves the interdomain linker of the ZP module itself - a region whose presentation within the filament completely depends on wrapping around two other protein molecules (Figure 2). A particularly relevant example are two O-glycosylation sites conserved within the ZP-N/ZP-C linker of mammalian ZP3 (Chalabi et al., 2006), the first one of which is also modified in chicken where it was shown to be important for sperm binding (Han et al., 2010). Projection of these sites onto the structure of polymeric UMOD suggests that site 1 is exposed at the end of α1β in the interdomain linker of ZP3, whereas site 2, which encompasses two or three closely spaced Ser/Thr residues depending on the species, is located within its β1. This periodically positions the O-glycosylation sites around the interface between ZP3 ZP-C and the ZP-N domain of the subsequent subunit (ZP1/2/4 in mammals or ZP1 in chicken), whose interaction most likely involves the invariant Trp of ZP3 that immediately follows site 1 and corresponds to V495 at the interface between UMOD ZP-C and ZP-N (Figure 7C, D). Remarkably, the same egg coat filament region is implicated by studies of gamete interactions in pig and bovine, where ZP subunit ZP4 plays a key role in fertilization by forming a sperm-binding complex together with ZP3 (Kanai et al., 2007; Yonezawa et al., 2012). The activity of this heterocomplex largely depends on tri- and tetra-antennary carbohydrate chains attached to pig ZP4 N220 (Yonezawa, 2014), and it is striking that this residue is predicted to be located just next to N271 - the only ZP3 sequon that carries the same two types of glycans (Yonezawa et al., 1999) - as well as in close proximity to ZP3 O-glycosylation sites 1 and 2 (Figure 7C). The latter may also contribute to sperm binding in the pig (Yurewicz et al., 1991), and - together with the near-invariant sequon of ZP3 at the beginning of α1β - additional glycosylation sites located in the same region may regulate the species-specificity of gamete interaction in other organisms.

Taken together, these considerations provide further support for the general idea that heteropolymeric ZP module protein filaments have the same basic architecture as UMOD; moreover, they strongly suggest that carbohydrate chains attached to adjacent ZP subunits wrapped by the ZP3 interdomain linker contribute to a common sperm-binding interface on the surface of egg coat filaments. Consistent with the aforementioned studies on pig and bovine ZP3/ZP4 (Kanai et al., 2007; Yonezawa et al., 2012), the difficulty to fully recapitulate such a complex system using non-interacting forms of ZP subunits expressed *in vitro* may have significantly hampered the decade-long search for a bona fide sperm counterpart of mouse and human ZP3.

### Interpretation of Pathologic Mutations in Human ZP Module Proteins

Because the conformations of the precursor and polymeric forms of ZP module-containing proteins are so different, the availability of structural information on how these molecules assemble into polymers will make it possible to more comprehensively assess the effect of missense mutations linked to human diseases.

For example, the structure of UMOD rationalizes the dominant negative effect of hearing loss-associated *TECTA* mutation Y1870C (Legan et al., 2005), which affects one of the two highly conserved Tyr residues in the ZP-N E’FG extension of inner ear protein α-tectorin, by showing that the corresponding residue in UMOD (Y402) is part of the interface with the ZP-C domain of the adjacent molecule (Figure 3B, right panel; Figure 3C, D). This suggests that the mutation disrupts the tectorial membrane not only by interfering with formation of the invariant C_1_-C_4_ disulfide of ZP-N (Monné et al., 2008), but also by directly affecting one of the polymerization interfaces of α-tectorin. Moreover, comparison of the location of this TECTA ZP-N mutation to that of UMOD ZP-N R415A (Figure 4) in light of the proposed mechanism of polymerization (Figure 5) explains why only the former has a dominant negative effect on polymerization.

A different but related example of dominant negative mutation is represented by recently described human ZP3 A134T, which causes female sterility due to empty follicle syndrome (Chen et al., 2017) and affects a residue at the N-terminal end of ZP-N βG. Due to the peripheral location of this strand in the ZP-N β-sandwich, the mutation is not expected to compromise the folding of the ZP3 precursor (Han et al., 2010; Monné et al., 2008); however, βG has a crucial role in polymerization by pairing with interdomain linker β1 from another subunit (right panel of Figure 3B). This clearly suggests that introduction of a polar side chain in βG will prevent the extension of the ZP-N hydrophobic core by β1, thus interfering with ZP filament formation and ultimately leading to lack of a ZP and oocyte degeneration.

### Biological Role of Proteins Containing Only the ZP-N Domain

Perhaps mimicking the sperm-binding ZP-N domain repeats found in the N-terminal region of egg coat proteins from both vertebrates and invertebrates, such as ZP2 and VERL (Callebaut et al., 2007; Monné et al., 2008; Raj et al., 2017), stacking of isolated ZP-N domains can generate filamentous structures *in vitro* (Jovine et al., 2006). However, in agreement with the invariant presence of both domains in the C-terminal region of all ZP module proteins (Bork and Sander, 1992; Litscher et al., 2015), the structure of polymeric UMOD conclusively shows that both ZP-N and ZP-C are essential for filament formation (Figure 2). This observation raises the question of what is the biological significance of the isolated ZP-N domains found in a number of non-egg coat proteins from worm to human (Jovine et al., 2006). Although further experimental studies will clearly be required to address this issue, our structural and mutagenesis data raise the intriguing possibility that these molecules may regulate the polymerization of full-ZP module proteins by attaching themselves to the growing end of their filaments. This is because, reminiscent of the pathogenic ZP-N mutations discussed in the previous section, such ZP-N-only subunits would effectively block further polymer extension due to the lack of a ZP-C domain.

### Potential Biotechnological Application of the UMOD Polymerization Scaffold

Consistent with the mosaic structure of ZP module-containing proteins (Bork and Sander, 1992; Litscher et al., 2015) and the filament architecture described in this work, functional studies on UMOD and egg coat proteins showed that the regions that precede the ZP module of these molecules can be replaced or fused with non-physiological domains without affecting protein incorporation into the respective filaments (Bokhove et al., 2016a; Jimenez-Movilla and Dean, 2011). Moreover, the unique structure of the UMOD ZP module polymer, where 24% of the surface area of each molecule is buried in interactions with four other subunits, makes it highly stable to proteolysis (Figure S1C-E) as well as remarkably resistant to chemical denaturation (Figure S7). These properties, which result from the significant reinforcement of subunit-subunit interactions by the continuous wrapping of interdomain linkers around the filament core (Figure 2), can be further strengthened by covalent cross-linking in other ZP module proteins, as observed for example in the case of egg coat subunits from fish to human (Greve and Wassarman, 1985; Nishimura et al., 2019; Yasumasu et al., 2010) or insoluble cuticlin components of the nematode larval alae (Sapio et al., 2005). These considerations explain why the ZP module has been evolutionarily selected for assembling a variety of protective matrices (Litscher et al., 2015). At the same time, together with the fact that UMOD polymers can be produced in recombinant form using cell lines that co-express hepsin (Bokhove et al., 2016a; Brunati et al., 2015), they raise the possibility of designing resilient Velcro-like biosurfaces decorated with different types and combinations of adhesive or catalytic domains.

## ACKNOWLEDGEMENTS

We thank the staff of the Swedish National Cryo-EM Facility, funded by the Knut and Alice Wallenberg Foundation, Family Erling Persson and Kempe Foundations, SciLifeLab, Stockholm University and Umeå University, for help with electron microscopes and pre-processing; the San Raffaele Advanced Light and Electron Microscopy BioImaging Center; T.C. Terwilliger for advice on density modification in PHENIX; T.I. Croll for help with ISOLDE; D.S. Goodsell for advice on figure generation using Illustrate and B. Forsberg for discussion. This work was supported by the Center for Innovative Medicine (Senior Investigator grant 2-537/2014 to L.J.), the Swedish Research Council (project grant 2016-03999 to L.J.), the Karolinska Institutet Research Foundation (grants 2016fobi50035 to L.J. and 2018-01646 to S.Z.-C.) and the Knut and Alice Wallenberg Foundation (project grant 2018.0042 to L.J.); the Ministry of Health, Singapore, NMRC grant (MOH-000382-00 to W.B.); the Italian Ministry of Health (RF-2010-2319394 and RF-2016-02362623 to L.R.).

## AUTHOR CONTRIBUTIONS

L.J. coordinated the study and designed the experiments together with L.R. and W.B.; A.S., S.Z.-C. and L.J. prepared UMOD material; A.S., S.Z.-C., M.C. and L.J. collected cryo-EM data; C.X., B.W., A.S. and L.J. performed structure determination, model building and refinement; M.B., C.S. and L.R. produced UMOD mutants and carried out immunofluorescence analysis; S.Z.-C., L.H. and A.S. expressed recombinant egg coat proteins and analyzed them by SEC-MALS; S.Y. performed fish egg coat digestions and mass spectrometry analysis; A.S. and L.J. wrote the manuscript with input from other authors.

## DECLARATION OF INTERESTS

The authors declare no competing interests.

## METHODS

### Purification and limited proteolysis of native human UMOD filaments

Full-length UMOD filaments (UMOD_fl_) were purified from the urine of one of the authors (L.J.) using the diatomaceous earth method (Serafini-Cessi et al., 1989), dialyzed against Milli-Q water and concentrated to 4 mg ml^-1^ using Amicon Ultra centrifugal filter units with Ultracel-50K (Merck Millipore). UMOD_e_ filaments were obtained by limited digestion of UMOD_fl_ with elastase (Jovine et al., 2002), dialysed overnight at 4°C against 10 mM Na-HEPES pH 7.0 and concentrated to 1 mg ml^-1^. Both UMOD_fl_ and UMOD_e_ samples were flash-frozen in liquid nitrogen and stored at −80°C until further use.

### Cryo-EM data collection

UMOD_fl_ (0.85 mg ml^-1^), UMOD_e_ (1 mg ml^-1^) and urine were applied in 3 µl-volumes onto glow-discharged Quantifoil Au R2/2 holey carbon 300 mesh grids (Quantifoil). Grids were blotted for 2.0 s and plunged into liquid ethane cooled by liquid nitrogen, using a Vitrobot Mark IV (Thermo Fisher Scientific).

All cryo-EM experiments were performed at the Cryo-EM Swedish National Facility, SciLifeLab, Stockholm. Movies for UMOD_fl_ or UMOD_e_ were collected with the EPU data acquisition software on a Titan Krios electron microscope (Thermo Fisher Scientific) operated at 300 kV, using a Gatan K2 Summit direct electron detector coupled with a Bioquantum energy filter with 20 eV slit. The defocus range for UMOD_fl_ was between −1.5 and −3.5 µm, the pixel size was 1.06 Å/pixel and the total dose was ∼40 electrons/Å^2^, distributed in 40 frames. For UMOD_e_, the defocus range was kept between −1.4 and −3.0 µm, the pixel size was 0.82 Å/pixel and the total dose was ∼45 electrons/Å^2^, distributed in 40 frames. Other data collection parameters are reported in Supplementary Table 1. Urine samples were imaged at 200 kV using a Talos Arctica microscope (Thermo Fisher Scientific) with a Falcon II detector (FEI).

Movie frames were aligned using MotionCor2 (Zheng et al., 2017) with dose compensation, as implemented in the Scipion on-the-fly processing pipeline (de la Rosa-Trevín et al., 2016).

### Helical reconstruction

CTFFIND (Rohou and Grigorieff, 2015) contrast transfer function (CTF) determination, particle picking, 2D classification, 3D classification and refinement procedures were performed using RELION (Zivanov et al., 2018).

For UMOD_fl_, a total of 2,300 micrographs were collected and analyzed (Figure S2). ∼24,000 filaments were manually picked and segments of 400 x 400 pixels, with 70 Å step size, were windowed out, yielding 412,322 particles for 2D classification. After two rounds of 2D classifications, ∼260,000 particles were chosen based on the appearance of 2D class images as well as particle image quality (rlnMaxValueProbDistribution) and resolution (rlnCtfMaxResolution) values estimated by RELION. Although most 2D classes had a distinct polarity, a few appeared to be symmetrical; the latter were thus further sub-classed in order to separate the opposite polarities. Multiple 2D classes with distinct image features were identified and at first we treated these 2D classes separately. Based on the power spectrum of the 2D class images, mathematically compatible helical rise values were calculated to be 125/n, with 1<=n<=10. After testing all these possible values, it was found that only with n=2 we could reproduce the angular views of any given 2D class, using the particles that belonged to it. However, reconstructed filaments originated from a single specific 2D class appeared to be completely flat in other angular views. This indicated that the distinct 2D class views might correspond to different angular views of a single type of filament with a helical rotation angle close to 180°. Thus, we pooled the 2D classes together and attempted helical reconstruction of the entire dataset. An *ab initio* low-resolution helical structure was generated using a featureless Gaussian cylinder with a diameter of 120 pixels and a length of 400 pixels as initial model. Preliminary 3D map reconstruction, however, suffered from severe flattening due to biased sampling of UMOD_fl_ filament at certain rotational angles. The fact that the helical rotation angle was close to 180° made it extremely challenging to piece together a reliable initial 3D model. Extensive topological analysis was thus carried out using a ∼30,000 particle subset that only included about one third of the over-represented front views, together with other angular views with stronger contrast (Figure S2). This smaller dataset, which had a better angular distribution, was used to both speed up the search and minimize the effect of the intrinsically biased sampling of filaments on the cryo-grids. We exhausted all the possible center value combinations of helical parameters within the ranges 160°-200° and 60-70 Å, using allowed windows of +/-5° and +/-2.5 Å for each run as well as 50% overlapping search ranges between different runs. Using the same featureless Gaussian cylinder as a starting model, all helical 3D reconstruction runs converged to ∼62.5 Å rise and ∼180.0° rotation, except in the cases where starting helical parameters and range were too far from these values (in such cases, the parameters simply stuck at the edge of the allowed window without converging). Further helical reconstruction runs with finer step sizes were thus subsequently performed to refine the 62.5 Å/180.0° values. If the output values of the searches agreed with the middle points of the corresponding allowed windows, we halved their permitted ranges and step sizes in the following run; if they did not agree, we shifted the middle points of the allowed ranges. During this process, we also gradually included additional 2D grouped particles that were not used during the initial helical parameter search (Figure S2). If the resulting helical 3D reconstruction runs failed to converge, we rolled back and included a smaller subset of new particles. Upon further refinement using +/-0.5° and +/-0.25 Å ranges, the parameters converged to a helical rise value of 62.5 Å and a helical rotation value of exactly 180.0°. After selecting the best pool of segments that shared this helical symmetry, these parameters were used for the 3D refinement of a set of 104,316 particles, which produced a 5.5 Å-resolution map of the complete filament segment. This served as a reference map (using information up to 7.0 Å resolution, in order to minimize overfitting) for additional direct 3D classification of all 412,322 particles, which captured more weakly contrasted particles with different angular views. Finally, 288,403 particles converged into a single 3D class that ultimately yielded significantly improved density after refinement with a smaller box size of 220 pixels, without the helical symmetry; according to a gold-standard 0.143-cutoff Fourier Shell Correlation (FSC) curve calculated using the PDBe FSC validation server (https://www.ebi.ac.uk/pdbe/emdb/validation/fsc), a resolution of 4.9 Å was reached for a central segment of density that includes five copies of the UMOD ZP module (with the majority of the filament core having a local resolution better than 4.2 Å, accordingly to ResMap (Kucukelbir et al., 2014)). A further improvement in map quality was obtained by density modification (DM) with PHENIX ResolveCryoEM (Terwilliger et al., 2020), which produced maps with estimated resolutions of 3.8 Å (multi-model-based DM)-4.0 Å (model-free DM; shown in Figure 1) that significantly facilitated model building by both enhancing general connectivity and resolving a large fraction of protein side chains.

Data processing of UMOD_e_ was performed similarly to UMOD_fl_, using 5,683 images to extract raw particles. Although UMOD_e_ is significantly more flexible and heterogeneous than UMOD_fl_, we managed to extract 252,438 usable particles (using a picking box of 350 x 350 pixels, with 70 Å step size) and, following a similar helical reconstruction protocol to that used for UMOD_fl_ data, we determined the helical twist and rise to be -179.9° and 62.7 Å, respectively. As in the case of UMOD_fl_, a direct 3D classification of all particles was then performed, which identified a set of 94,937 homogenous particles that converged into a single 3D class. 3D refinement of these particles, using a 280 Å box, fine angular sampling and no helical symmetry, produced a 6.0 Å-resolution map (FSC = 0.143 criterion). Finally, postprocessing of the latter with ResolveCryoEM yielded densities with estimated resolutions of 3.9 or 4.1 Å, depending on whether a multi-model representation was used or not for DM, respectively.

Cryo-EM maps have been deposited in the Electron Microscopy Data Bank (EMDB) with accession codes EMD-10553 (UMOD_fl_) and EMD-10554 (UMOD_e_).

### Model building, refinement, validation and analysis

Coordinates of human UMOD EGF IV/ZP-N domains (residues T296-L429) and ZP-C domain (excluding linker β-strand 1 and the EHP-containing C-terminal propeptide; residues P466-F587) were extracted from chain A of the X-ray crystallographic model of the polymerization-inhibited UMOD precursor (PDB ID 4WRN) (Bokhove et al., 2016a). UCSF Chimera (Pettersen et al., 2004) was used to first place into the central portion of the UMOD_fl_ cryo-EM map a copy of the EGF IV/ZP-N fragment, whose position - despite less defined density for the relatively flexible EGF domain - was unequivocally indicated by comparison of the UMOD_fl_ and UMOD_e_ densities and consistent with a clearly corresponding elongated region of the map (EGF IV/ZP-N(I); correlation 0.82). Subsequently, a copy of the ZP-C domain model was docked into the remaining part of the central region of the map (ZP-C(I); correlation 0.88). Additional copies of ZP-C and EGF IV/ZP-N (ZP-C(II) and EGF IV/ZP-N(II)) were then added adjacent to the previously placed EGF IV/ZP-N(I) and ZP-C(I) models, respectively, by taking into account helical symmetry information. At this stage, it became evident that an uninterrupted (and unaccounted for) extended stretch of density contacting both ZP-N(I) and ZP-C(I) linked the C-terminus of ZP-N (II) to the N-terminus of ZP-C (II). The latter domains, together with EGF IV(II) connected to ZP-N (II), were therefore assigned to a single molecule of UMOD (chain A, corresponding to UMOD 3 in Figure 2). Based on symmetry considerations, we also concluded that ZP-C(I) and EGF IV/ZP-N(I) belong to two distinct additional copies of UMOD (chains B and C, respectively; corresponding to UMOD 2 and UMOD 4 in Figure 2), both of which interact with chain A as well as with each other. The resulting initial set of coordinates, consisting of one chain encompassing the whole polymerization region of UMOD (A) and two half chains (B, C), was subjected to molecular dynamics (MD) flexible fitting in Namdinator (Kidmose et al., 2019) and manually rebuilt using Coot (Casañal et al., 2019) as implemented in CCP-EM (Burnley et al., 2017). After further improvement by Cryo_fit (Kim et al., 2019b), as well as additional rebuilding in Coot (whose carbohydrate-building tool (Emsley and Crispin, 2018) was used to add the N-glycan chains attached to EGF IV N322, ZP-N N396 and ZP-C N513) and ISOLDE (Croll, 2018), the model was real-space refined against the UMOD_fl_ data in PHENIX (Afonine et al., 2018a) using a data/restraint weight of 0.8, a non-bonded weight of 250.0 and restraints generated using the starting coordinates as a reference. Protein and carbohydrate coordinates were validated with PHENIX (Afonine et al., 2018b)/MolProbity (Williams et al., 2018) and CCP4’s Privateer (Agirre et al., 2015), respectively; model-to-map validation was carried out with EMRinger (Barad et al., 2015). Since taken together they represent all the protein-protein interactions found in the UMOD polymer, all three UMOD chains have been included in the final deposited model, which consists of 592 protein residues (V293-F587 (chain A); V443-F587 (chain B); V293-S444 (chain C)) and 14 N-glycan residues.

The model of elastase-treated UMOD was generated by first rigid-body-fitting the atomic coordinates of the UMOD_fl_ ZP module regions and then further optimizing the position of the EGF IV domains of chains A and C, which - consistent with the flexibility observed in the crystal structure of its precursor form (Bokhove et al., 2016a) - in UMOD_e_ are rotated by ∼15° about the Z axis with the filament model in front view. After deletion of C-terminal residues disordered due to lack of an interacting D8C domain, refinement and validation were performed essentially as described for UMOD_fl_; the final model consists of 588 protein residues (S292-T585 (chain A); V443-S583 (chain B); V292-S444 (chain C)) and 18 N-glycan residues.

Homology modelling of the UMOD EGF I-III domain region was performed with the I-TASSER threading server (Yang and Zhang, 2015), which produced a set of coordinates that was consistent with the conserved disulfide bond pattern of other EGF domains (1-3, 2-4, 5-6) (Wouters et al., 2005). I-TASSER was also used to generate models of the central region of UMOD (residues E149-S291, including the D8C domain) in parallel with *ab initio* modelling using the Robetta server (Kim et al., 2004). The top model produced by the latter, which had closely positioned pairs of Cys (consistent with the suggestion that all Cys in UMOD are engaged in disulfide bonds (Friedmann and Johnson, 1966; Hamlin and Fish, 1977)) and exposed to the solvent the side chains of glycosylated residues N232 and N275 (van Rooijen et al., 1999), was selected and refined by MD simulation in YASARA Structure (Krieger et al., 2009). Models combining the refined coordinates of EGV IV/ZP module with either D8C or the whole N-terminal region of UMOD were generated by first fusing in Coot molecular fragments fitted into the EM density and then energy minimizing the resulting coordinate sets with YASARA.

Secondary structure was assigned using STRIDE (Frishman and Argos, 1995); protein-protein interfaces were analyzed using PISA (Krissinel and Henrick, 2007), PIC (Tina et al., 2007) and the AnalyseComplex command of FoldX (Delgado et al., 2019); evolutionary conservation was assessed with ConSurf (Ashkenazy et al., 2016). Structural figures were generated with PyMOL (Schrödinger, LLC), UCSF Chimera (Pettersen et al., 2004)/ChimeraX (Goddard et al., 2018) and Illustrate (Goodsell et al., 2019); unless specified, they were based on the refined coordinates of UMOD_fl_.

Atomic coordinates have been deposited in the Protein Data Bank (PDB) with accession codes 6TQK (UMOD_fl_) and 6TQL (UMOD_e_).

### Sequence-structure analysis

Hidden Markov model logos were generated with Skylign (Wheeler et al., 2014), using as input the Pfam (El-Gebali et al., 2019) seed alignment for the Zona_pellucida family (PF00100), modified to include the sequence of human UMOD instead of its rat homologue and manually edited to correct a misalignment of UMOD C_b_ to the last conserved Cys of the ZP module (C_8_). Homology models of mouse ZP3 ZP-C and pig ZP4 ZP-N were generated using MODELLER (Webb and Sali, 2016), based on sequence-structure alignments produced by HHpred (Zimmermann et al., 2018).

### DNA constructs

Wild-type or truncated UMOD cDNA constructs were cloned in pcDNA3.1(+) (Thermo Fisher Scientific) and HA- or FLAG-tags were inserted after the signal peptide, between T26 and S27 in the protein sequence (Schaeffer et al., 2009). Mutation of the hepsin cleavage site (586-RFRS-589/AAAA; 4A mutant) was described previously (Schaeffer et al., 2012); constructs pcDNA3.1(+)/UMOD-CCS or pcDNA3.1(+)/UMOD-EHP expressed proteins truncated at residues S589 or S614, respectively. Polymerization interface mutants were generated using a QuikChange Lightning mutagenesis kit (Agilent) following the manufacturer’s instructions; primers were designed using the QuikChange Primer Design server (https://www.agilent.com/store/primerDesign Program.jsp).

Co-expression of FLAG- and HA-tagged UMOD isoforms was performed using a bicistronic vector pVITRO-hygro-mcs (Invivogen). For this purpose, the sequence of FLAG-tagged wt UMOD was subcloned from pcDNA_FLAG-UMOD into the EcoRV site of pVITRO-hygro-mcs, downstream of the mouse elongation factor-1α promoter; the sequences of HA-tagged UMOD isoforms (wt or mutants) were subcloned from pcDNA_HA-UMOD in the Bst1107I site, downstream of the rat elongation factor-1α promoter of the same bicistronic vector.

For mammalian expression of medaka ZI-1,2 (P223-Q591) and ZI-3 (Y74-V420) constructs, synthetic genes (ATUM) were subcloned into vector pHLsec3 (Raj et al., 2017), in frame with a sequence encoding a C-terminal 6His-tag.

All DNA constructs were verified by DNA sequencing (Eurofins Genomics) before transfection.

### Recombinant protein expression and purification

MDCK cells were grown in Dulbecco’s Modified Eagle’s Medium (DMEM) supplemented with 10% fetal bovine serum, 200 U ml^-1^ penicillin, 200 μg ml^-1^ streptomycin and 2 mM glutamine at 37°C, 5% CO_2_. For characterization of UMOD mutants, stable cell populations were generated by transfection with Lipofectamine 2000 (Thermofisher) following the manufacturer’s protocol. Selection was started 24 h after transfection by adding 0.5 mg/ml G418 (Thermofisher) and was pursued for 1-2 weeks in order to obtain a population of G418-resistant cells.

For analyzing the oligomerization state of egg coat protein precursors, medaka ZI-1,2 (which does not contain N-glycosylation sites) was expressed in HEK293T cells (DuBridge et al., 1987), grown in DMEM medium supplemented with 4 mM L-Gln 2% fetal bovine serum and transiently transfected using 25 kDa branched PEI (Aricescu et al., 2006; Bokhove et al., 2016b). GnTI-deficient HEK293S cells (Reeves et al., 2002) were used to express ZI-3 carrying Endoglycosidase H (Endo H)-sensitive Man5GlcNAc2 N-glycans. These carbohydrate chains were then enzymatically trimmed to single GlcNAc residues during protein purification, which was performed by batch immobilized metal ion affinity (IMAC) using nickel agarose slurry (Ni-NTA, QIAGEN or Ni Sepharose High Performance, GE Healthcare) and size-exclusion chromatography (SEC) using a Superdex 200 Increase 10/300 GL column (GE Healthcare) equilibrated against 20 mM HEPES pH 7.5, 150 mM NaCl (Bokhove et al., 2016b).

### Immunoblot

Cell lysis, medium precipitation and western blot experiments were performed essentially as described (Schaeffer et al., 2009), using mouse purified anti-HA.11 Epitope Tag monoclonal (clone 16B12) (Biolegend 901502; 1:1,000 dilution), mouse anti-β-actin monoclonal (clone AC-74) (Sigma-Aldrich A2228; 1:20,000 dilution) and rabbit affinity-isolated anti-FLAG polyclonal (Sigma-Aldrich F7425; 1:1,000 dilution).

### Immunofluorescence

Cells grown on coverslips were fixed in 4% paraformaldehyde for 15 min, permeabilised for 10 min with 0.5% triton when indicated and blocked for 30 min with 10% donkey serum. Cells were labelled for 1 h 30 min with the indicated primary antibodies at room temperature, followed by 1 h incubation with the appropriate Alexa-Fluor conjugated secondary antibodies (Thermo Fisher Scientific; 1:500). They were then stained for 5 min with 4,6-diamidino-2-phenylindole (DAPI) and mounted using fluorescence mounting medium (DAKO, Agilent). Pictures were taken with an UltraVIEW ERS spinning disk confocal microscope (Zeiss 63X/1.4, UltraVIEW ERS-Imaging Suite Software; PerkinElmer) or a DM 5000B fluorescence upright microscope (Leica DFC480 camera, Leica DFC Twain Software; Leica Microsystems). All images were imported in Photoshop CS (Adobe) and adjusted for brightness and contrast.

The antibodies used for these experiments were mouse purified anti-HA.11 Epitope Tag monoclonal (clone 16B12) (Biolegend 901502; 1:500 dilution), rabbit affinity-isolated anti-FLAG polyclonal (Sigma-Aldrich F7425; 1:500 dilution), rat Anti-HA High Affinity monoclonal (clone 3F10) (Roche 11867423001; 1:500 dilution), goat affinity purified anti-c-Myc polyclonal (Novus Biologicals NB 600-335; 1:500 dilution) and mouse anti-KDEL monoclonal (clone 10C3) (Enzo ADI-SPA-827; 1:200 dilution).

### Fish egg coat digest preparation

Unfertilized eggs, isolated from spawning female Japanese rice fish following procedures approved by the ethics committee of Sophia University (approval number 2016-006), were crushed in 50 mM Tris-HCl pH 7.5-buffered saline containing 5 mM ethylenediaminetetraacetic acid and 5 mM iodoacetic acid. After centrifugation at 5,000 rpm for 30 s, the supernatant was decanted. This procedure was repeated several times to completely remove yolk protein and cell debris. Isolated egg coats were used for digestion experiments, using HCE and LCE purified from crude hatching liquid (Yasumasu et al., 1989a, 1989b). Fifty UFEs were incubated for 1 h at 30 °C in 100 ml 50 mM Tris-HCl pH 8.0 containing either 2.3 µg purified HCE or 1.2 µg purified LCE, or their mixture. The resulting material was fractionated using a HiLoad 16/60 Superdex 200 GL column (GE Healthcare) equilibrated with phosphate-buffered saline (PBS; 20 mM phosphate buffer pH 7.2, 150 mM NaCl). Two eluted protein peaks were collected and separately re-chromatographed using the same column. Final single peaks were analysed by SDS-PAGE on 12.5% gels, as well as by native PAGE using 5-20% gradient gels.

### Cross-linking analysis of the egg envelope digest

1.75 ml egg envelope digest (∼100 µg ml^-1^ in PBS) was mixed with 50 µl 0.1 mg ml^-1^ disuccinimidyl suberate (DSS). Cross-linking was allowed to proceed for 20 min at 25 °C and quenched with the addition of 100 µl 1 M Tris-HCl pH 8.0. The protein solution was then concentrated using an Amicon Ultra 15 Ultracel-10K device (Millipore). Samples were desalted using a Millipore Ziptip C18, concentrated and analyzed with a MALDI-TOF MS AXIMA-Performance mass spectrometer (Shimadzu). The matrix consisting of 10 mg 3,5-dimethoxy-4-hydrocinnamic acid was dissolved in a 1 ml reaction mixture containing 50% (*v/v*) acetonitrile, 0.05% (*v/v*) trifluoroacetic acid.

### Mass spectrometric analysis of the subunit composition of the egg coat digest

TOF-MS analysis of SEC-purified F2 (Figure 4A), a sample containing three protein fragments of approximately 37, 18 and 16 kDa (Figure 4B, C), showed two sharp peaks and two broad peaks (Figure 4D). Whereas the m/z values of the former (18706.26 and 17846.13) closely matches the predicted molecular weights of ZP-C- and ZP-N-containing fragments of the ZI-1,2 subunit (18704 (S388-G557) and 17852 (T221-D387)), the latter (m/z 36559.54 and 39801.56) was assigned to a digestion product of ZI-3 that contains both ZP-N and ZP-C (Y74-T393, 34817 kDa) and is heterogeneously glycosylated at N184. The identity of the fragments was confirmed by Edman degradation, and a similar pattern was obtained from the analysis of SEC-purified F1 (Figure 4E).

To determine the subunit composition of F2 and F1, a cross-linking approach was employed. Each SEC peak was cross-linked by DSS, an amine-reactive compound that covalently links lysine residues, and their molecular weights were determined by MS. Cross-linking of F2 produced a single peak (Figure 4F), whose m/z 74029 matches the sum of the molecular weights of the 37, 18 and 16 kDa digestion products (73111.93-76353.95). Considering that F2 migrates as a single band on native-PAGE (Figure 4C), this result suggests that F2 is a heterotrimeric complex containing a single copy of each digestion product. Unlike F2, DSS-cross-linked F1 consisted of several regularly spaced species, whose m/z values differed by multiples of ∼18000-20000 (Figure 4G); this observation immediately suggests that the cross-linked complexes reflect a repeated subunit structure within the egg coat filaments. Considering that the two digestion fragments of ZI-1,2 (corresponding to the two halves of the ZP module) have an average molecular weight of ∼18.3 kDa and taking into account that the species in F1 have higher molecular weights than the complex in F2 by native-PAGE (Figure 4C), the MS peaks with m/z values 55605.79 and 768038.12 can be assigned to incomplete cross-linking products consisting of half ZI-1,2 + ZI-3 and two copies of ZI-3, respectively. Similarly, higher m/z peaks can be interpreted as follows: 96658,72 = half ZI-1,2 + 2x ZI-3; 114825.53 = 2x half ZI-1,2 + 2x ZI-3; 133329.12 = half ZI-1,2 + 3x ZI-3; 152398.32 = 2x half ZI-1,2 + 3x ZI-3 (Figure 4G). These assignments are consistent with the difference in intensity between the ZI-1,2 18 and 16 kDa digestion products in F1 and those in F2 (Figure 4B), and - considering that F1:F2 ratio is ∼0.8 (Figure 4A and (Iuchi and Yamagami, 1976)) - indicate that heteromeric interactions (ZI-1,2/ZI-3) are more abundant than homomeric ones (ZI-3/ZI-3) in the UFE HCE/LCE-digest.

### SEC-MALS analysis

Ettan LC high-performance liquid chromatography system equipped with UV-900 detector (Amersham Pharmacia Biotech; λ= 280 nM), coupled with miniDawn Treos MALS detector (Wyatt Technology; λ= 658 nM) and an Optilab T-rEX dRI detector (Wyatt Technology; λ= 660 nM) was used to analyse the absolute molar mass of ZI proteins. Separation was performed at a flow rate of 0.5 ml min^-1^, using a Superdex 200 Increase 10/300 GL column (GE Healthcare) equilibrated against 20 mM HEPES pH 7.5, 150 mM NaCl. Data processing and weight-averaged molecular mass calculations were done with Astra software (Wyatt Technology Corporation). SEC-MALS experiments were repeated independently twice, with each experiment including *de novo* expression, purification and measurement of each protein, rather than just a repeated analysis of the same material. All measurements of each protein agreed, but for clarity the results of a single experiment are presented in Figure S6.

### Depolymerization assays

Native UMOD filaments, purified as described above, were diluted to 0.25 mg ml^−1^ in 0-8 M urea, 1 mM EDTA and incubated overnight at 37°C before centrifugation at 110,000 g in a Beckman TLA 100 rotor for 1 h at 10°C. The top half of the supernatant of each sample was carefully removed for analysis and, after discarding the remaining solution, pellets were solubilized in SDS sample buffer. Samples were analysed on 10% SDS–PAGE gels.

---

**Figure S1.**
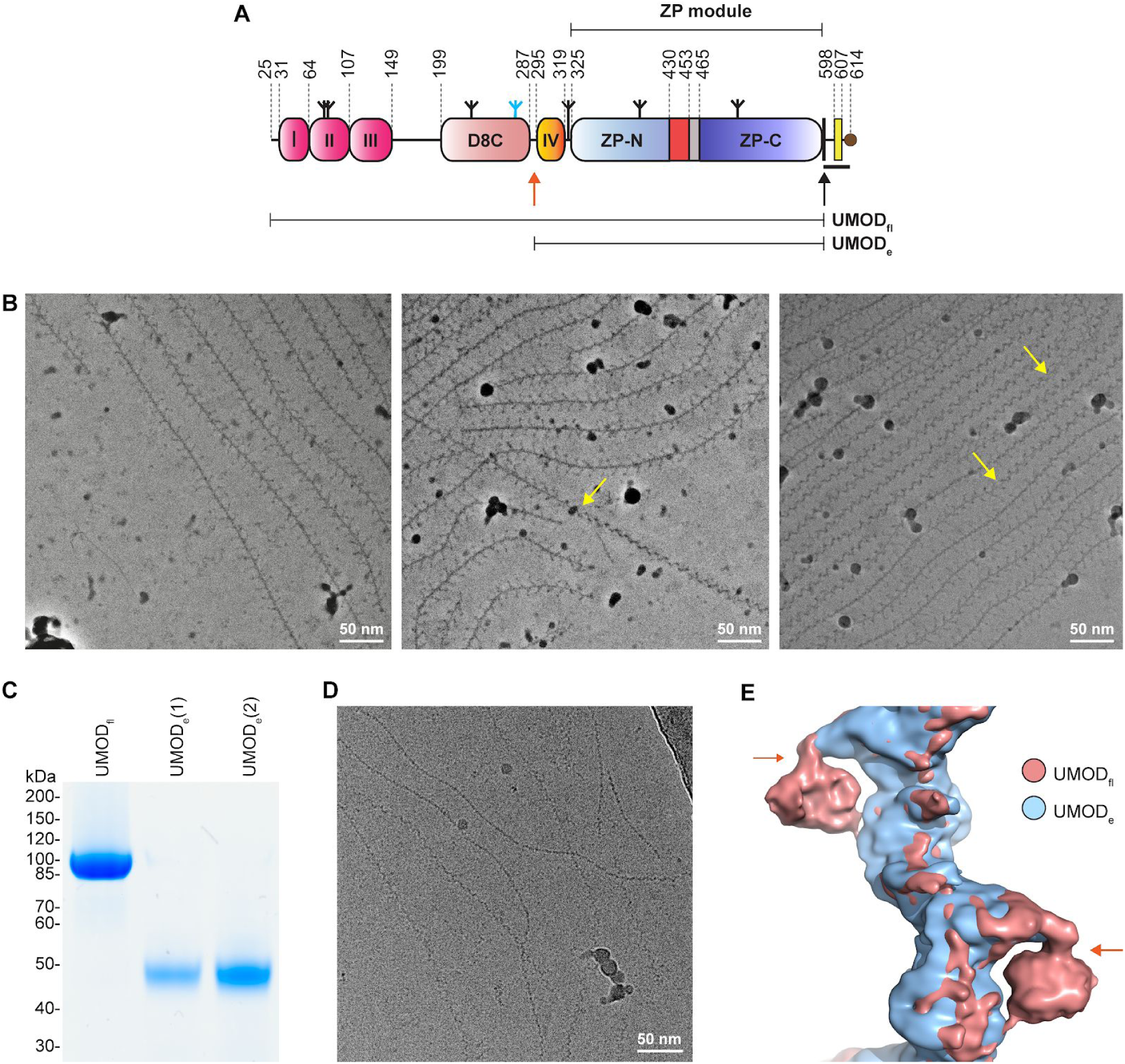
Comparison of Full-Length and Elastase-Treated UMOD Filaments. (A) Domain organisation of the secreted human UMOD precursor. Magenta, EGF I-III; salmon, D8C domain; orange, EGF IV; light blue and deep blue, ZP-N and ZP-C domains; red, ZP-N/ZP-C linker; grey, internal hydrophobic patch (IHP); black, CCS; yellow, EHP. A thick black horizontal line marks the CTP, with a brown circle depicting the GPI anchor attachment. Inverted tripods show N-glycans, with the high-mannose chain attached to D8C N275 coloured cyan. Black and orange arrows indicate the position of the hepsin (F587|R588) and elastase (S291|S292) cleavage sites, respectively, with thin horizontal bars indicating the extent of UMOD_fl_ and UMOD_e_. (**B**) Representative Volta phase plate micrographs of native UMOD_fl_ filaments. Although tree/front views are predominant, a number of zig-zag/side views can be seen in the right-most micrograph. The yellow arrows show examples of how twisting of individual UMOD filaments generates both views. (**C**) Reducing Coomassie-stained SDS–PAGE analysis of the UMOD_fl_ (6 µg; lane 1) and UMOD_e_ (3 and 5 µg; lanes 2, 3) material used for structure determination. (**D**) Representative micrograph of UMOD_e_ filaments, showing the absence of branches. (**E**) Superposition of the UMOD_fl_ (salmon) and UMOD_e_ (blue) cryo-EM maps shows that only the former shows density for a globular domain protruding from the core of the filaments. This reveals the approximate location of the elastase cleavage site, corresponding to the N-terminus of UMOD_e_, within the structure of UMOD_fl_ (orange arrows).

**Figure S2.**
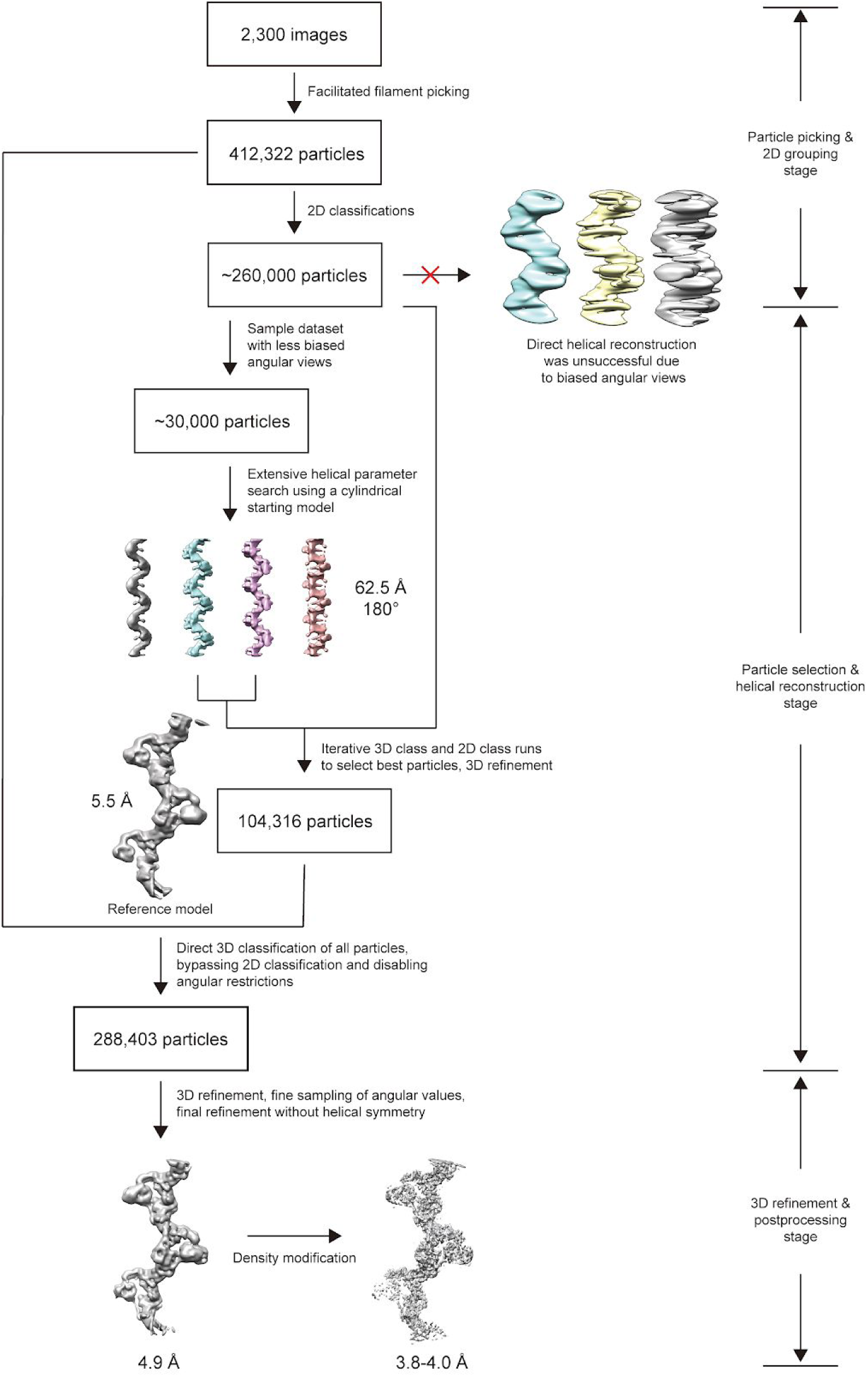
3D Helical Reconstruction of UMOD Filaments. Helical reconstruction of the 3D density map of UMOD_fl_ was performed in three steps. (1) Particle picking and 2D grouping. Filaments were picked with the help of contrast-enhancing programs and pre-filtered based on contrast and clarity of the segments. Once helical pitch and power spectrum were obtained, we attempted direct 3D helical reconstruction, but failed to obtain reasonable density. (2) Searching for the true helical parameters and selecting the best group of particles. After realizing that it was necessary to pool different 2D classes together, for time efficiency we selected a subset of 30,000 particles with the best contrast and better angular distribution and used them to iteratively sample potential combinations of helical parameters consistent with the 2D views. Once the correct helical parameters were determined, we gradually included more particles in the 3D classification and refinement procedures. Since some of the angular views were intrinsically weaker than the others, we bypassed the 2D classification step and proceeded with 3D classification of all the extracted particles. Eventually, we selected 288,403 particles that aligned with a single 3D class density. (3) 3D refinement and postprocessing. All selected particles were combined and used in a few final rounds of 3D refinement, with gradually reduced sampling steps. This eventually led to a 3D helical reconstruction density of UMOD_fl_ with significantly improved map features and nominal resolution, which was further enhanced by density modification.

**Figure S3.**
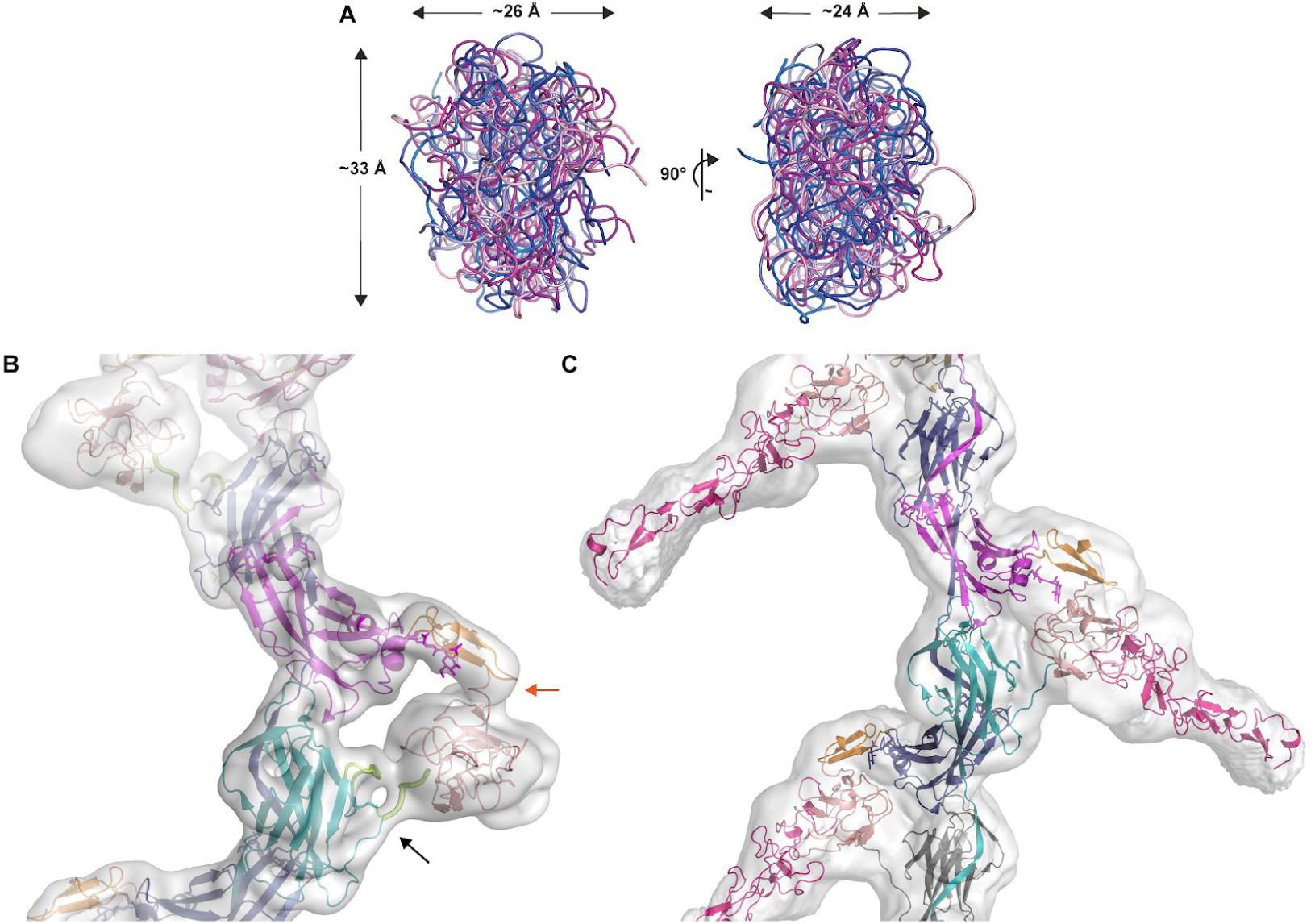
Docking of D8C and EGF I-III Domain Models into the Density Map of UMOD_fl_. (A) D8C domain models created using I-TASSER (Yang and Zhang, 2015) (blue shades) or Robetta (Kim et al., 2004) (magenta shades) have approximately the same overall dimensions. (**B**) Consistent with the location of the elastase cleavage site (orange arrow), which immediately precedes the EGF IV domain (orange), the top D8C model generated by Robetta (salmon) can be straightforwardly docked into the globular density protruding from the core of the filament. The unmodified cryo-EM map of UMOD_fl_ is shown, and a black arrow indicates the C_6_527-C_8_582 disulfide, which orients the C-terminal tail of mature UMOD (thick lemon tube) towards D8C. The latter also packs against the loop that connects C_6_527 to βD (thin lemon tube). (**C**) A density map contoured at a level approximately 4.5 times lower than in panel B shows elongated features extending from the D8C domain, which are consistent with the expected dimensions of UMOD EGF I-III (pink).

**Figure S4.**
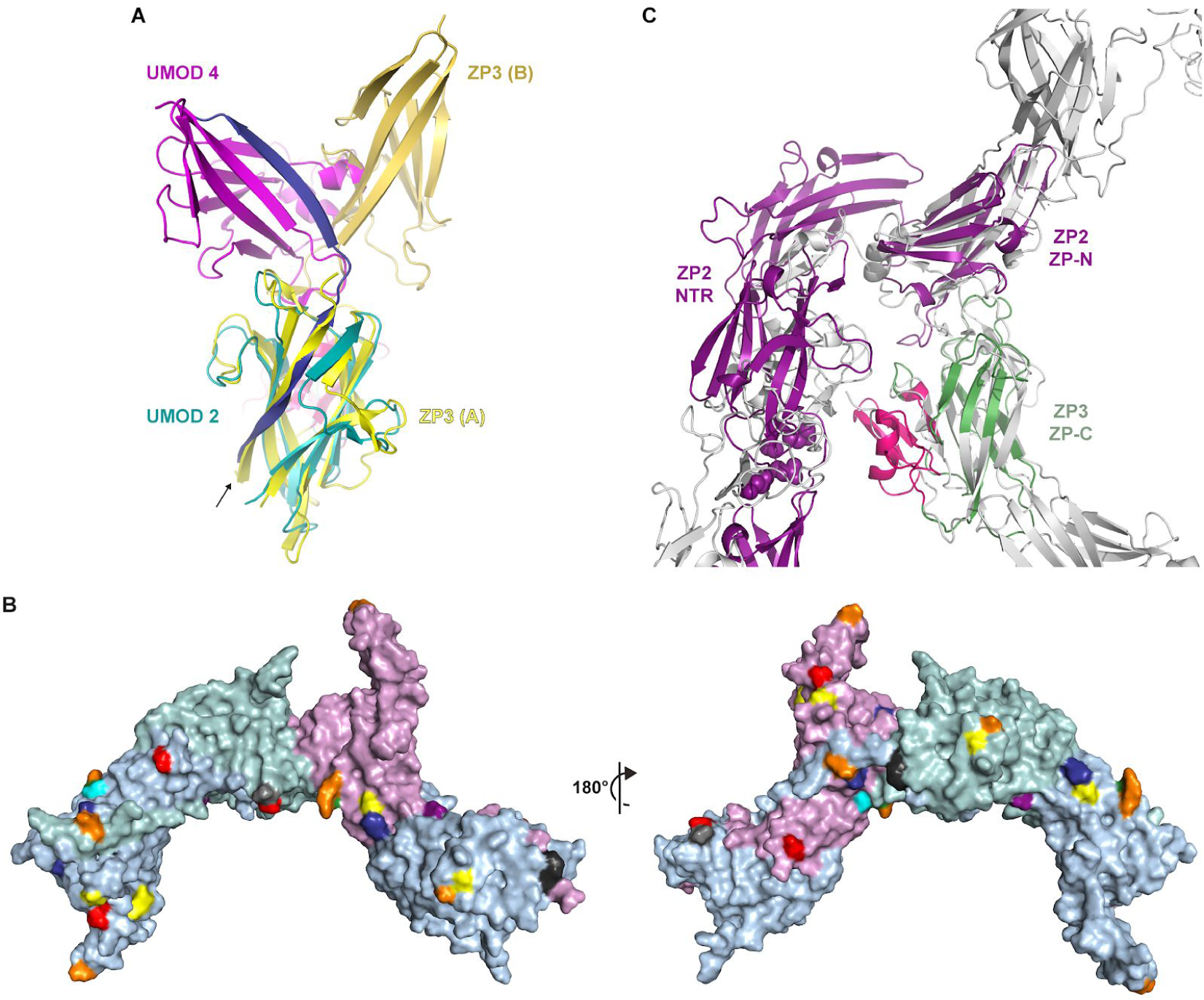
UMOD Subunit Interactions: Comparison with the Homodimeric cZP3 Precursor and Consistency with Features of Other ZP Module Proteins. (**A**) Superposition of the ZP-C domains of a polymeric UMOD subunit (UMOD 2 of Figure 2; teal) and one of the two subunits of the cZP3 precursor (Han et al., 2010) (chain A of PDB 3NK3; yellow, with the ZP3-specific subdomain colored hot pink). The ZP-N counterparts (magenta and orange-yellow, respectively) that interact with these domains are differently positioned relatively to the corresponding ZP-Cs, although both interfaces are formed by the same elements. The arrow indicates how superimposing the ZP-C domains brings the EHP of ZP3 in the same position as α1β in the interdomain linker of the UMOD subunit that follows UMOD 2 (UMOD 3; blue). (**B**) The ZP module interface observed in the UMOD filament is compatible with the expected solvent exposure of the N- and O-glycosylation sites of other ZP module proteins. Predicted N-glycosylation sites of human glycoprotein 2 (orange), α- and β-tectorin (yellow and blue), ZP2 (green), ZP3 (grey), ZP4 (cyan), chicken ZPD (purple), as well as O-glycosylation site 1 of chicken ZP3 (black), are mapped onto the surface of three adjacent UMOD chains (A, light blue; B, light teal; C light magenta) based on sequence-structure alignments. UMOD N-glycosylation sites are shown in red. (**C**) Homology models of the N-terminal repeat region (NTR) of mouse ZP2 plus its ZP-N domain (Monné et al., 2008) (purple) and the ZP-C domain of mouse ZP3 (green) were superimposed on the ZP-N and ZP-C domains of two adjacent UMOD filament subunits (grey), respectively; subsequently, ZP2 NTR was approximately oriented like the N-terminal branch of the same UMOD subunit used for the ZP-N/ZP-N superposition. The resulting model shows that, akin to UMOD EGF I-III/D8C, the ZP-N domain repeats that precede the ZP module of ZP2 can project from the core of egg coat filaments without interfering with subunit polymerization interfaces. Similarly, the C-terminal subdomain specific to ZP3 (hot pink) is predicted to be positioned laterally to the egg coat filament body and potentially face the ovastacin cleavage site in the second N-terminal repeat of ZP2 (spheres).

**Figure S5.**
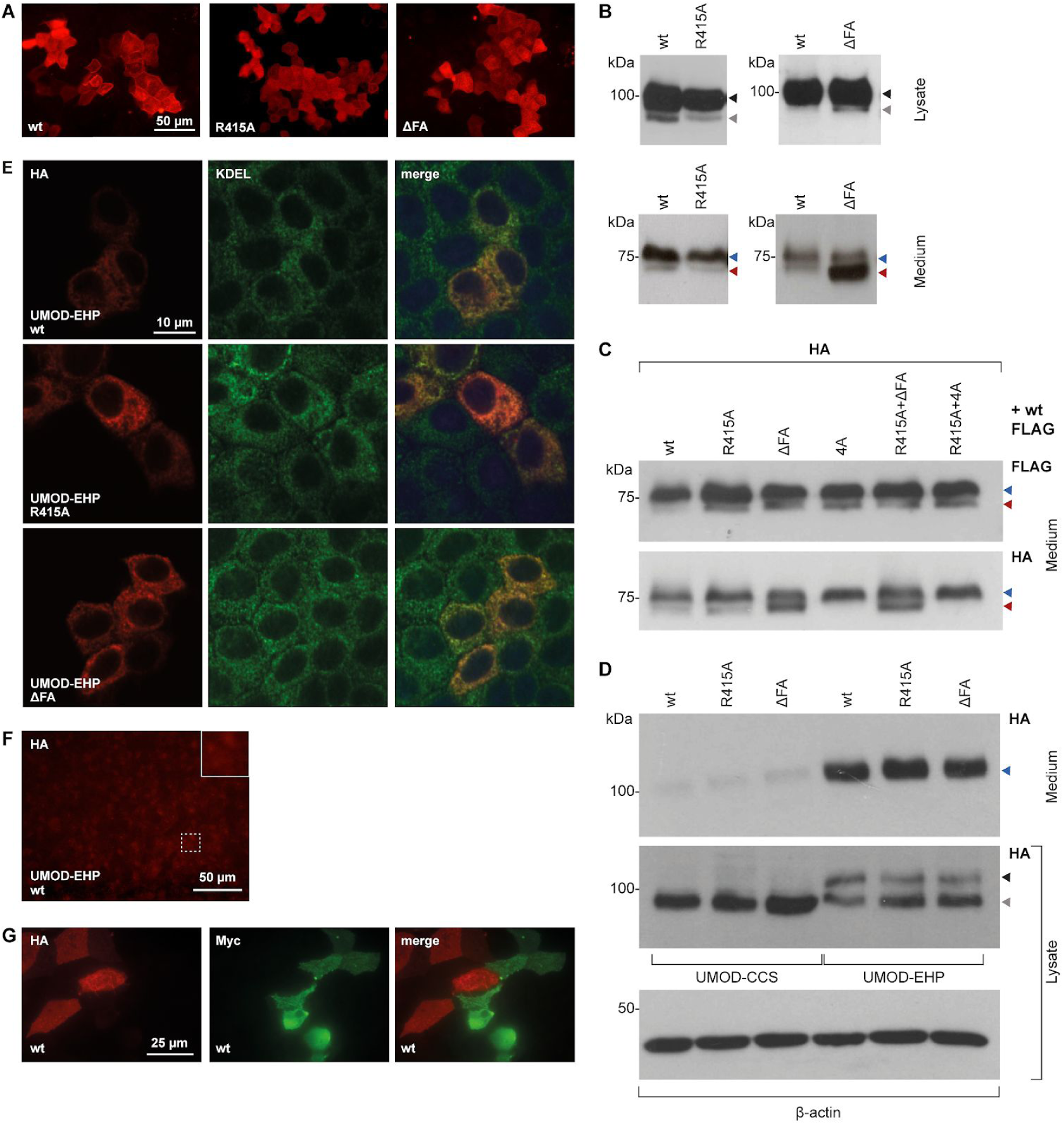
Analysis of the Effects of UMOD Mutations on Protein Expression, Secretion and Filament Formation. (**A**) Immunofluorescence of permeabilized MDCK cells expressing full-length wt UMOD or mutants R415A or ΔFA. No intracellular aggregation is observed. (**B**) MDCK cells stably expressing R415A or ΔFA mutants of full-length UMOD. Immunoblots of total cell lysates (top panels) indicate that both wt protein and polymerization mutants are mainly present as a fully glycosylated isoform (upper band, black arrow), in addition to a minor ER-glycosylated species (lower band, grey arrow). Immunoblot analysis of PNGase F-deglycosylated proteins secreted by MDCK cells (bottom panels) shows that neither the R415A nor the ΔFA mutation affects protein secretion in the culturing medium. Blue arrows indicate proteins cleaved within the juxtamembrane region between GPI and EHP; red arrow mark proteins that were processed at the CCS. These results demonstrate comparable intracellular trafficking and secretion of wt and mutant isoforms. Note that the ΔFA mutation increases the amount of protein that is processed at the CSS, suggesting that alteration of the FXF motif affects the accessibility of the closely located cleavage site (Figure 3B, left panel); despite this, filaments are completely absent in the case of the mutant (Figure 4A, right panel), further underlying the specific effect of the mutation on UMOD polymerization. (**C**) Immunoblot of deglycosylated proteins released in the culturing medium of MDCK cells stably co-expressing FLAG-tagged wt UMOD and HA-tagged wt or mutant isoforms. Co-expression of mutant UMOD does not alter the cleavage of the FLAG-tagged wt protein, suggesting that the dominant negative effect of the ΔFA and 4A mutants is not caused by abnormal processing of wt UMOD. (**D**) Immunoblot of UMOD in the cell lysate and conditioned medium of MDCK cells transiently transfected with the indicated HA-tagged isoforms. The presence of the EHP motif is required for efficient protein exit from the ER, as suggested by comparing the intracellular levels and secretion of UMOD-CCS and UMOD-EHP. (**E**) MDCK cells transiently expressing ZP-N (R415A) and ZP-C (ΔFA) mutants of UMOD-EHP. Immunofluorescence of permeabilized cells shows absence of intracellular polymers in both wt and mutant forms. (**F**) Immunofluorescence analysis of unpermeabilized MDCK cells expressing UMOD-EHP. Lack of membrane-anchoring prevents localization and polymerization of the protein at the plasma membrane. (**G**) Co-culture of MDCK cells stably expressing HA-tagged (red) or Myc-tagged (green) wt UMOD. Filaments are uniformly colored, suggesting that polymerization depends on incorporation of membrane-bound monomers instead of cleaved monomers released in the culture medium.

**Figure S6.**
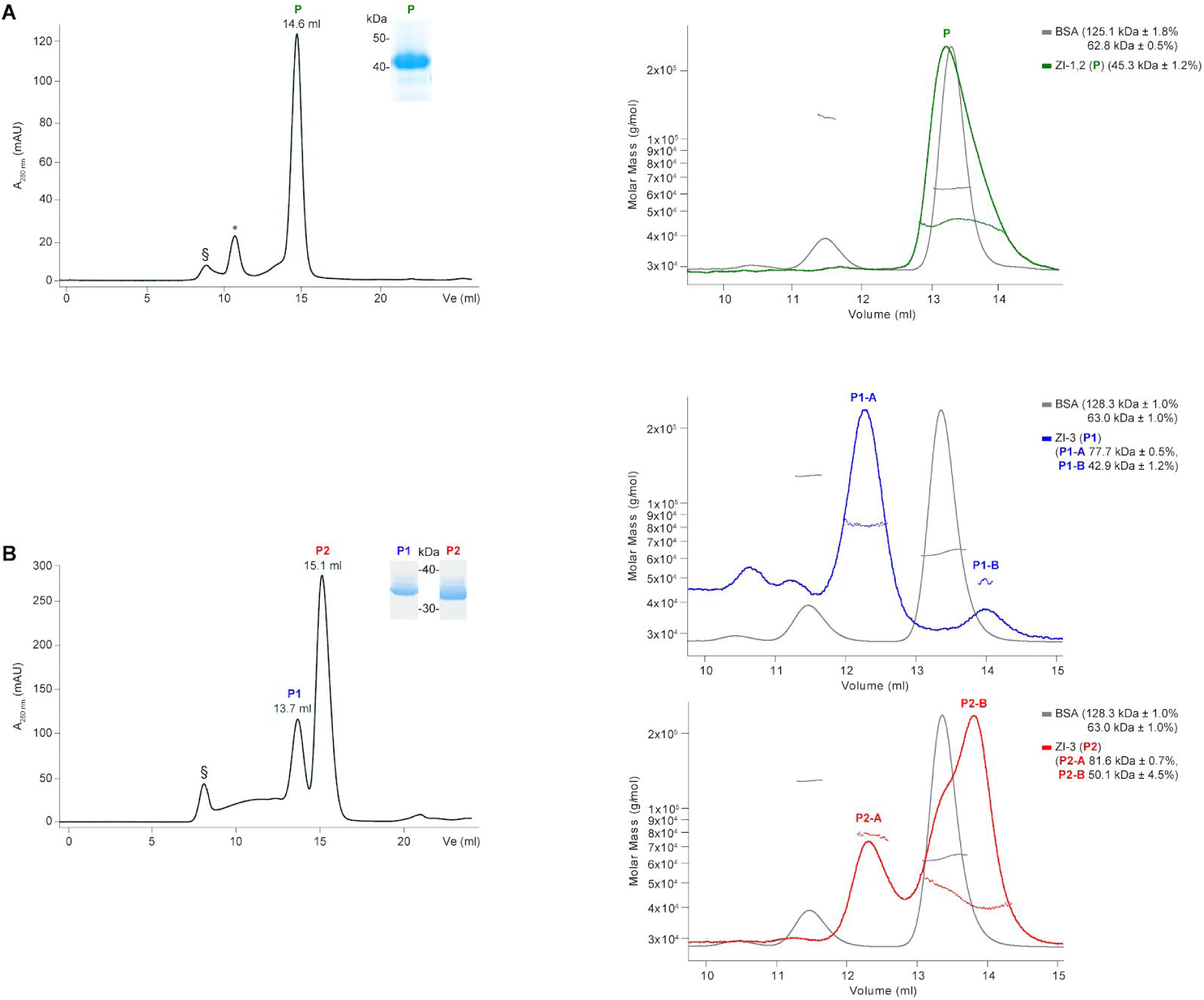
Analysis of the Oligomeric State of Fish Egg Coat Proteins. IMAC-captured His-tagged proteins were purified by SEC (left panels) and relevant elution peaks were analyzed by reducing Coomassie-stained SDS-PAGE (left panel insets) and SEC-MALS (right panels). (**A**) Medaka ZI-1,2 is a monomer in solution. (**B**) Medaka ZI-3 elutes as two SEC peaks that correspond to homodimeric (P1) and monomeric (P2) forms of the glycoprotein. SEC void volume and contaminant peaks are indicated by § and *, respectively. SEC-MALS experiments were calibrated using bovine serum albumin (BSA).

**Figure S7.**
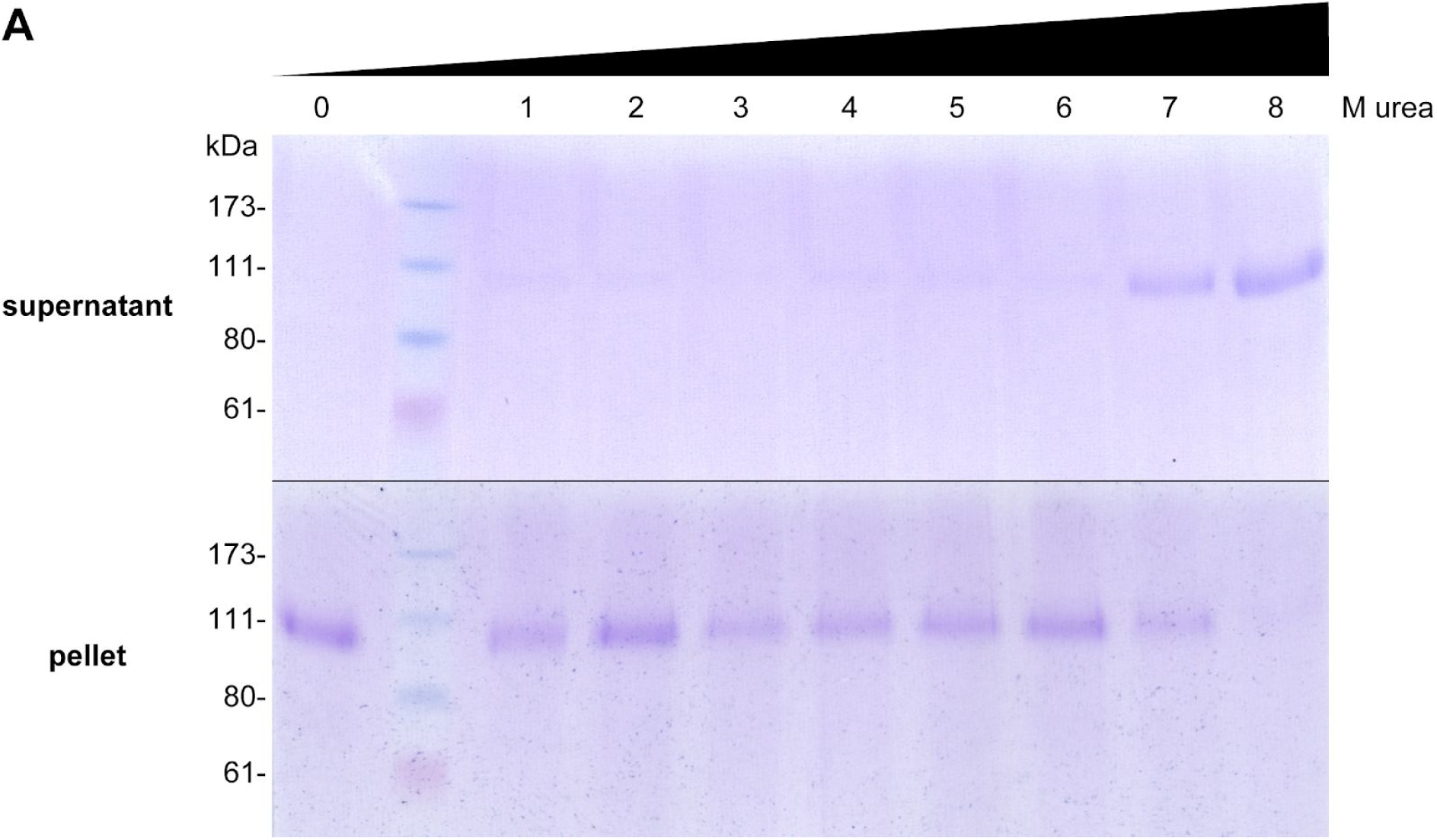
UMOD Filaments are Stable in 6 M Urea. Coomassie-stained SDS-PAGE analysis of supernatant and pellet fractions of purified native UMOD filaments, incubated with increasing amounts of urea. No significant breakdown of the polymers is observed at urea concentrations below 7 M.

**Table S1.** Cryo-EM data collection, refinement and validation statistics.

|  | **UMOD<sub>fl</sub>**<br>EMDB: EMD-10553<br>PDB: 6TQK | **UMOD<sub>e</sub>**<br>EMDB: EMD-10554<br>PDB: 6TQL |
| --- | --- | --- |
| **Data collection and processing** |  |  |
| Magnification | 130,000 | 165,000 |
| Voltage (kV) | 300 | 300 |
| Electron exposure (e<sup>-</sup>/Å<sup>2</sup>) | 39.6 | 44.6 |
| Defocus range (μm) | -1.5 – -3.5 | -1.4 – -3.0 |
| Pixel size (Å) | 1.06 | 0.82 |
| Symmetry imposed | Helical (62.5 Å rise, 180.0° rotational angle) | Helical (62.7 Å rise, -179.9° rotational angle) |
| Initial particle images | 412,322 | 252,438 |
| Final particle images | 288,403 | 94,937 |
| Map resolution (Å)<br>(FSC = 0.143) | 4.9 | 6.0 |
| Map resolution range (Å) | 3.8 – 5.6 | 4.6 – 6.8 |
| DM map resolution (Å)<br>(FSC<sub>ref</sub> = 0.5; multi-model / model-free) | 3.8 / 4.0 | 3.9 / 4.1 |
| **Refinement** |  |  |
| Initial model used (PDB code) | 4WRN | 6TQK |
| Model composition |  |  |
| Non-hydrogen atoms | 4,814 | 4,812 |
| Protein residues | 592 | 588 |
| Carbohydrate residues | 14 | 18 |
| *B* factors (Å<sup>2</sup>) |  |  |
| Protein | 116 | 115 |
| Ligand | 130 | 135 |
| R.m.s. deviations |  |  |
| Bond lengths (Å) | 0.003 | 0.003 |
| Bond angles (°) | 0.481 | 0.575 |
| **Validation** |  |  |
| MolProbity score | 1.13 | 1.48 |
| Clashscore | 1.59 | 3.30 |
| Rotamer outliers (%) | 0 | 0 |
| Ramachandran plot |  |  |
| Overall Z-score (RMSD) | 1.30 (0.31) | -1.59 (0.33) |
| Favored (%) | 96.6 | 94.9 |
| Allowed (%) | 3.4 | 5.1 |
| Disallowed (%) | 0 | 0 |
| Model-to-data fit |  |  |
| CC(mask) | 0.7353 | 0.7153 |
| CC(volume) | 0.7319 | 0.7073 |
| CC(peaks) | 0.4156 | 0.3870 |
| EMRinger score | 2.37 | 1.64 |

## Notes

### Competing Interest Statement

The authors have declared no competing interest.

